# *BABYBOOM*-like expression in the cowpea egg and central cell enables parthenogenesis, endosperm development, and viable haploid seed formation

**DOI:** 10.64898/2026.02.08.704694

**Authors:** Itzel Amasende-Morales, Osvaldo Ruiz-Maciel, Gloria Leon-Martínez, Joann Conner, Sailaja Bhogireddy, Quetzely Ortiz-Vasquez, Huanan Su, Juana de la Cruz, Judith Lua, Karen Ramírez, Miguel Vallebueno-Estrada, Stefano Bencivenga, Ramona Hartmann, Nial Gursanscky, Matteo Riboni, Martina Juranic, Melanie L. Hand, Susan D. Johnson, Brett Ferguson, Ueli Grossniklaus, Peggy Ozias-Akins, Anna M.G. Koltunow, Jean-Philippe Vielle-Calzada

## Abstract

Parthenogenesis or fertilization-independent embryogenesis occurs at low frequency in sexual plants. Expression of *BABYBOOM-like* (*BBML*) and *PARTHENOGENESIS* (*PAR*) genes in the egg cell of several diploid dicot crops induce parthenogenesis at varying frequency; however, recovery of viable haploid seeds has rarely been reported, perhaps due to a lack of viable endosperm formation. In the legume cowpea (*Vigna unguiculata* L. Walp), ectopic egg cell expression of the endogenous *BBML* homolog (*VuBBML1*) and *PAR* from *Taraxacum officinale* induces parthenogenesis; however, seeds abort as endosperm formation is blocked following self-pollination. Expression of *VuBBML1* in both the egg cell and central cell, together with central cell fertilization following self-pollination, results in viable seeds that germinate and give rise to haploid plants. *VuBBML1* has a functional role in the formation of cowpea embryo and endosperm seed compartments. This finding opens possibilities for establishing double haploid production during homozygous parental breeding, and asexual seed induction for fixing hybrid vigor in cowpea.

## Main Text

Most flowering plants rely on sexual reproduction for seed formation. Meiosis is necessary for the development of male and female gametes. Within the female gametophyte, double fertilization of the egg cell and central cell by two sister sperm cells generates the embryo and endosperm, while the surrounding ovule tissues give rise to the maternal seed coat. Parthenogenesis refers to the autonomous development of embryos in the absence of fertilization of the egg cell, giving rise to haploid embryos carrying the gametic chromosome number of n instead of 2n.

The sporadic formation of haploid embryos by parthenogenesis has been reported in at least 74 species belonging to 21 flowering plant families [1]; however, it rarely leads to germinable seed formation and haploid plants. The development of nutritive endosperm is presumably needed via central cell fertilization for viable haploid seeds to form. Yet spontaneous parthenogenesis and viable haploid seed formation in the absence of endosperm formation was recently reported in a specific sunflower genotype suggesting that other nutritional pathways can support embryo development in this eudicot species [2]. Parthenogenesis also occurs in plants that can form seeds asexually by apomixis. These plants also avoid meiosis in female gamete formation and can develop a functional endosperm giving rise to clonal seeds [3]. The recovery of haploid seed is beneficial for crop breeding as the initial step of double haploid production, reducing the time and cost necessary to generate homozygous parental lines [4,5].

Parthenogenesis has been successfully induced in monocots, including pearl millet, rice, maize, and sorghum following ectopic egg cell expression of the *AINTEGUMENTA-LIKE APETALA2/ETHYLENE RESPONSE FACTOR* (*AP2/ERF*) domain transcription factor *PsASGR-BBML* from the gametophytic apomict *Cenchrus squamulatus* (syn. *Pennisetum squamulatum*) [6–8]. Similarly, rice and maize homologs *OsBBM1* and *ZmBBM1* also trigger efficient parthenogenesis when ectopically expressed in the egg cell [9,10]. Parthenogenetic induction has been deployed together with meiosis avoidance to generate synthetic apomixis in monocot crops such as rice and sorghum [8, 11–14]. By contrast, *in vivo* ectopic expression of *BBML*-like genes in the wild-type egg cell of eudicots such as tetraploid tobacco and diploid Arabidopsis has resulted in viable progeny at frequencies lower than 1% [15,16]. In lettuce, the K2-2 zinc finger class gene, *PARTHENOGENESIS* (*PAR*) from apomictic *Taraxacum officinale* (*To*; dandelion) can also induce egg cell division in the absence of fertilization; however, recovery of haploid seeds has not been reported [17].

Cowpea (*Vigna unguiculata* L. Walp; 2n= 22) is a subsistence legume crop domesticated in sub-Saharan Africa that provides calories and protein for millions of smallholder farmers [18]. As part of a program aiming to enable smallholders to save and sow hybrid seeds through synthetic apomixis, the induction of viable parthenogenetic cowpea seed was examined using the *Taraxacum officinale PAR* gene (*ToPAR*) gene and the cowpea *PsASGR-BBML* homolog. Prior developmental analyses in the cowpea variety IT86D-1010 established a developmental calendar correlating stages of male and female gametogenesis with floral development [18]. Here, early embryo and endosperm formation were cytologically examined following controlled pollinations. In 97% of IT86D-1010 ovules (n=166), syncytial endosperm formation initiated in the central cell prior to cellular division of the zygote (Figure S1 and Table S1). Only 3% of the cowpea ovules showed early zygote division prior to endosperm initiation, suggesting endosperm initiation in cowpea IT86D-1010 typically precedes embryogenesis following double fertilization.

Variety IT86D-1010 was transformed with a construct containing the native *ToPAR* promoter or the Arabidopsis *pAtRKD2* egg cell specific promoter driving the *ToPAR* cDNA (*pToPAR::ToPAR* and *pAtRKD2::ToPAR* constructs, respectively; Figure 1a). Four of 24 *pToPAR:ToPAR* T0 transformants analyzed three days after emasculation (DAE) showed embryo-like bodies containing up to 10 nuclei with rarely visible cell walls, at frequencies ranging between 1.25% and 3.65% (Figures 1b and 1c; Tables 1 and S2). These frequencies are within range of those reported for parthenogenesis in other eudicot species [15,16]. Ten of 18 *pAtRKD2:ToPAR* T0 transformants also displayed parthenogenetic embryo initiation at frequencies ranging between 10 and 20% (Figure S2), with pods producing lower-than-expected number of seeds. In all cases the central cell remained quiescent, and free nuclear divisions characteristic of initial stages of endosperm development were not observed. In emasculated non-transgenic plants, the female gametophyte degraded or remained intact with no autonomous egg cell divisions at three DAE. Self-pollinated *pToPAR:ToPAR* transformants only produced sexually derived diploid seed. Crosses of female *pAtRKD2::ToPAR* and *pToPAR:ToPAR* transformants with male non-transgenic plants or transformants homozygous for a molecular marker construct (*pEF1A::DsRed-Express*) showed that the *pAtRKD2::ToPAR* and *pToPAR:ToPAR* transgenes were not transmitted through the female gametophyte, suggesting that all embryo sacs expressing *ToPAR* in the egg cell aborted before reaching seed maturity (Table S3).

**Figure 1.**
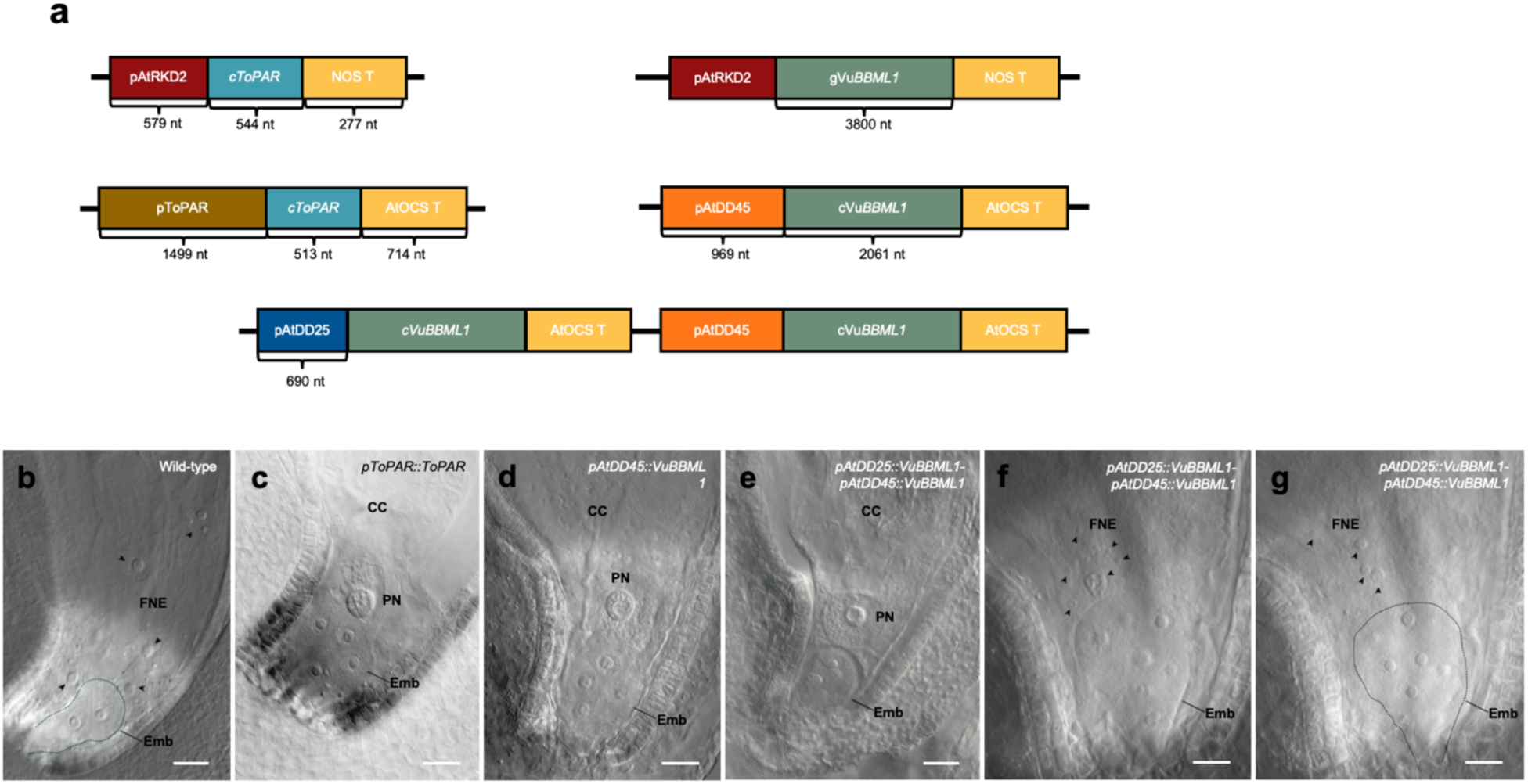
Parthenogenesis in cowpea after ectopic expression of *ToPAR* or *VuBBML1* in specific cells of the female gametophyte. **(a)** Wild-type cowpea plants were transformed with different constructs harboring the *PAR* gene of *Taraxacum officinale* under its native promoter; or with the cowpea *VuBBML1* gene under the Arabidopsis *pAtRKD2*, *pAtDD25*, and *pAtDD45* promoters. **(b)** Two-cellular asymmetric embryo and free nuclear endosperm in a wild-type ovule following pollination and double fertilization; free nuclei are marked by arrowheads. **(c)** Ovule of a *pToPAR::ToPAR* transformant three days after emasculation (DAE); a parthenogenetic embryo is visible in the absence of endosperm development. **(d)** Ovule of a *pAtDD45::VuBBML1* transformant two DAE; a parthenogenetic embryo is visible in the absence of endosperm development. **(e)** Ovule of a *pAtDD25::VuBBML1-pAtDD45::VuBBML1* transformant three DAE; a two-cellular asymmetric embryo is visible prior to endosperm development. **(f)** and **(g)** Ovule of a *pAtDD25::VuBBML1-pAtDD45::VuBBML1* transformant three DAE (two focal planes); a large multicellular embryo develops in synchrony with free nuclear divisions of the endosperm (arrowheads). Abbreviations: Emb, embryo; FNE, free nuclear endosperm; PN, polar nucleus. Scale bar: 10μm.

A candidate cowpea *PsASGR-BBML* homolog (*VuBBML1*) was initially identified from genomic sequences of varieties IT97K-499-35 and IT86D-1010 [19], using the assembled IT97K-499-35 genome as a reference [20]. Of 25 cowpea genes containing two predicted *AP2/ERF* domains, only *VuBBML1* clustered with *BBML* genes functionally shown to induce parthenogenesis when expressed in the egg cell (Figure S3a). qRT-PCR expression showed that *VuBBML1* is preferentially expressed in cowpea ovules at the zygote stage rather than in other organs (Figure S3b). The 4.442 kb genomic sequence of the *VuBBML1* candidate from IT97K-499-35 is identical to that found in IT86D-1010 [21] and is composed of a 218 bp 5’ untranslated region (UTR), 9 exons (8 introns) and a 422 bp 3’ UTR (Figure S4).

Because Arabidopsis promoters *pAtRKD2* and *pAtDD45* drive expression in the egg cell of transgenic cowpea (Figure S5) [22, 23], both were independently fused to the *VuBBML1* genomic sequence to test for parthenogenesis induction at one to three DAE (*pAtRKD2::VuBBML1* and *pAtDD45::VuBBML1* constructs, respectively; Figure 1a). The *pAtRKD2::VuBBML1* construct was introduced into the IT86D-1010 background, and the *pAtDD45::VuBBML1* construct was introduced into a hybrid cowpea genotype (*Hyb18*) resulting from a cross between varieties IT86D-1010 and Tvu1503. Cytological analyses of untransformed IT86D-1010 and *Hyb18* emasculated flowers showed absence of parthenogenesis (n=150 and n=121, respectively). By contrast, 13 out of 26 transgenic lines containing the *pAtRKD2::VuBBML* construct developed parthenogenetic embryos at frequencies comprised between 8 and 25% (Figure S6). Crosses of *pAtRKD2::VuBBML1* transformants with four different cowpea varieties confirmed that parthenogenesis is transmitted and maintained at similar rates in the resulting hybrid cowpea backgrounds (Table S4). A total of six out of ten emasculated T0 *pAtDD45::VuBBML1* transformants showed parthenogenetic embryo initiation at a frequency of 5.5% (Figure 1d; Tables 1 and S5).

The embryonic-like bodies formed in *pAtRKD2::VuBBML1* and *pAtDD45::VuBBML1* parthenogenetic lines resembled those formed in *pAtRKD2::ToPAR* and *pToPAR::ToPAR* transformants, containing up to 10 nuclei with rarely defined cells walls. This suggested potential defects in cellularization and pattern formation during parthenogenesis induction. Nuclear divisions were not observed in the central cell of any of these transformants. Emasculated flowers aborted three to five DAE without producing seeds, indicating that the parthenogenetically derived embryos are unable to progress beyond a few divisions. Both *pAtRKD2::VuBBML1* and *pAtDD45::VuBBML1* transformants showed fertility defects associated with parthenogenesis, and reciprocal crosses to untransformed plants confirmed that in both cases the transgenic construct was only transmitted through male gametophytes, suggesting that all developing seeds expressing *VuBBML1* in the egg cell aborted before reaching maturity (Table S6). Consequently, all T0 transformants only gave rise to sexually derived diploid plants originating from fertilized egg cells. Collectively, these results demonstrated that the expression of *VuBBML1* in the cowpea egg cell can induce parthenogenesis but viable haploid seeds are not generated following self-fertilization of female gametophytes containing parthenogenetic embryos.

Given the lack of maternal transmission within parthenogenetic plants harboring the *pAtRKD2::ToPAR* or *pAtRKD2::VuBBML1* constructs, pollen tube attraction to the synergids of ovules containing these constructs was examined. A male homozygous transgenic line containing a *DsRed-Express* fluorescent marker driven by a constitutively expressed soybean promoter (*pGmEF1a::DsRed-Express*) was crossed to heterozygous *pAtRKD2::ToPAR* or *pAtRKD2::VuBBML1* transgenic plants showing parthenogensis. In both cases approximately 50% of ovules that did not inherit the construct causing parthenogenesis developed sexually derived embryos expressing *DsRed-Express*. By contrast, 55 to 60% of the ovules containing the construct causing parthenogenesis did not show endosperm initiation; however, they showed *DsRed-Express* expression within synergids but not within the adjacent parthenogenetic embryo (Table S7 and Figure S7), suggesting that pollen tubes can penetrate the synergids of transgenic female gametophytes but are unable to trigger endosperm formation. However, endosperm development is never initiated.

In self-pollinated flowers of Arabidopsis, *BBM* is expressed in the embryo and transiently in the primary endosperm nucleus at initial stages of endosperm development [16]. Mutant analyses suggested a role for Arabidopsis *BBM* in endosperm proliferation and cellularization [16]. In cowpea, the Arabidopsis *pAtDD25* promoter drives reporter expression to the central cell (Figure S5). Given that cowpea female gametophytes containing parthenogenetic embryos did not initiate endosperm development under self-pollinating conditions, we hypothesized that the induction of autonomous divisions of the central cell might sustain subsequent haploid seed progression. To test this possibility, the Arabidopsis *pAtDD25* promoter was used to express *VuBBML1* in the central cell prior to double fertilization. *Hyb18* wild-type plants were transformed with a dual *VuBBML1* construct containing two *VuBBML1* genes: one driven by the *pAtDD45* promoter to target egg cell expression, and the other driven by the *pAtDD25* promoter to target central cell expression (*pAtDD25::VuBBML1-pAtDD45::VuBBML1;* Figure 1a).

Cytological analyses of all 19 emasculated transgenic T0 plants showed parthenogenetic embryos at frequencies ranging between 8.2% and 31.8% (Tables 1 and S5). Strikingly, 11 out of the 19 transformants also showed autonomous proliferation of free nuclei in the central cell at frequencies ranging between 2.1% and 9.4%, indicating that *VuBBML1* can induce free nuclear proliferation of the central cell in the absence of fertilization (Tables 1 and S5; Figures 1d to 1f). Contrary to self-pollinated untransformed plants where endosperm development generally initiates prior to embryogenesis (Figure S1), *pAtDD25::VuBBML1-pAtDD45::VuBBML1* transformants initiated embryogenesis prior to autonomous nuclear divisions in the central cell (Figure 1e). Remarkably, the first division of the egg cell in the initiating parthenogenetic embryo was asymmetric (Figure 1e), giving rise to a small apical cell and larger basal cell, a pattern equivalent to the first zygotic division in sexually derived cowpea embryos following egg cell fertilization (Figure 1b). This pattern of parthenogenetic embryo initiation was not observed in *pToPAR::ToPAR* and *pDD45::VuBBML1* transformants that targeted *VuBBML1* expression only to the egg cell.

In contrast to *pAtDD45::VuBBML1* plants that were semi-sterile (Tables 1 and S8), pods arising after self-pollination of T1 *pAtDD25::VuBBML1-pAtDD45::VuBBML1* transformants contained 12 to 14 seeds per pod, similar to untransformed plants (Figure 2a; Tables 1 and S8). However, a mixture of seed phenotypes were observed, including seeds equivalent in size to those of control untransformed plants (average length: 0.82±0.06 cm; average width: 0.53±0.05 cm; n=20), smaller seeds with a smooth or shriveled seed coat (Fig 2a and 2b; average length: 0.41±0.05 cm; average width: 0.33±0.03 cm; n=20), and flattened or collapsed aborted seeds that did not germinate (Figure 2b). The smaller seeds (smooth or shriveled) were slow to germinate *in vitro* compared with untransformed controls (untransformed average: 1.6±0.5 days, n=30; small seeds: 14.2±6.3 days, n=53). Their seedlings often showed early developmental defects that delayed or hindered their developmental progression (Figure 2c). The resulting plantlets were dwarfed, yellowish and did not reach flowering (Figure 2d).

**Figure 2.**
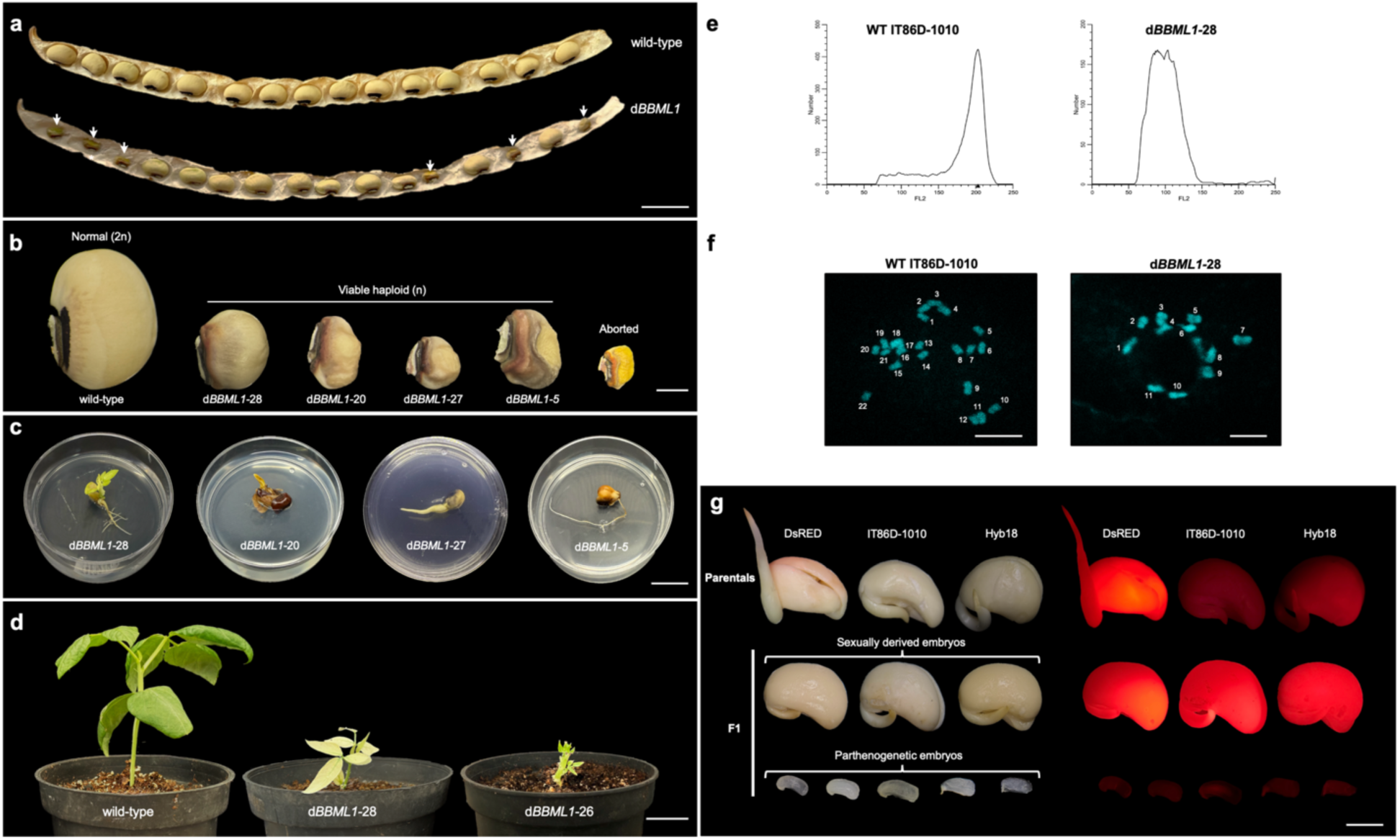
Viable parthenogenesis in cowpea after simultaneous expression of *VuBBML1* in the egg cell and central cell. (**a**) Seed set in mature pods of wild-type and *pAtDD25::VuBBML1-pAtDD45::VuBBML1* transgenic individuals (*dBBML1*); small haploid seeds are marked by arrows. (**b**) Normal and small smooth or shriveled seeds produced after self-pollination of a dBBML1 transformant; aborted seeds are lenticular or collapsed. (**c**) Examples of normal and defective seedlings generated after *in vitro* germination of small haploid seeds; acronyms correspond to seeds shown in (**b**). (**d**) Phenotypes of haploid plantlets under greenhouse conditions. (**e**) Ploidy assessment of nuclei from parthenogenic haploid (*dBBML1-28*) and control (WT IT86D-1010) diploid leaves. (**f**) Chromosome counts in parthenogenetic haploid (*dBBML1-28*) and control (WT IT86D-1010) diploid root tips. (**g**) Fluorescent phenotypes of transgenic F1 isolated embryos generated after crossing a *pAtDD25::VuBBML1-pAtDD45::VuBBML1* transgenic plant (as female) to a constitutive *DsRed-Express* marker (as male). A weak autofluorescent signal is present in non-transgenic female parental backgrounds IT86D-1010 and Hyb18. Strong red fluorescence is present in embryos isolated from normal size F1 diploid seeds but absent in embryos isolated from F1 small haploid seeds. Scale bars: (a) = 1.5 cm; (b) = 0.2 cm; (c) = 1 cm; (d) = 2 cm; (f) = 10 μm; (g) = 0.5 cm.

Ploidy analyses of leaves by flow cytometry (Figure 2e), and cytogenetic analysis of root cells (Figure 2f), confirmed that plants recovered from the smaller seeds were haploid. Maternally derived haploid plants are expected to appear homozygous at all loci, particularly at alleles showing heterozygosity in the hybrid maternal background. Genotyping T1 plants using more than 1.9 million single nucleotide variants (SNVs) showing heterozygosity in T0 *Hyb18* transformants indicated that the T1 plants derived from small seeds showed only a single nucleotide variant across all 11 cowpea chromosomes, confirming their maternal haploid nature (Table S9; Supplemental File 1). In T1 plants homozygous for the transgene, the overall fertility and frequency of the small seed class was increased, in comparison to T0 hemizygous transformants (on average 18.8% vs 12.7% of small seeds; with a maximum of 40% in *dBBL1-700* T1 progeny; Tables 1 and S8). This suggested that viable parthenogenesis is dosage sensitive and dependent on the level of *VuBBML1* expression.

As *pAtDD25::VuBBML1-pAtDD45::VuBBML1* dual promoter expression is controlled at the gametophytic (n) level and given that the egg cell and central cell are both derived from a functional meiosis, a maximum rate of 50% parthenogenesis was expected in single copy T0 transformants hemizygous for the transgenic construct. Since the maximum proportion of ovules containing parthenogenetic embryos in self-pollinated plants was 31.8%, the frequency of transgene transmission through male and female gametes was estimated by conducting reciprocal crosses with the *pGmEF1a::DsRed-Express* transgenic line. Observation of embryos dissected from mature seeds derived from these crosses showed that single copy hemizygous *pAtDD25::VuBBML1-pAtDD45::VuBBML1* transformants were transmitted through either male of female gametes at nearly 100% rates (Table 2), This suggested that pollen grains carrying the transgene are able to germinate and fertilize at similar rates as wild-type pollen, and that female gametophytes carrying the construct but not initiating parthenogenesis give rise to sexually derived seeds following double fertilization. Embryos isolated from either small smooth or shriveled seeds showed a similar organization with partially developed cotyledons covering an embryonary axis that contained a radicle without obvious defects (Figure 2g). The absence of *DsRed-Express* expression in these small embryos confirmed their maternal nature, demonstrating that they were derived via parthenogenetic development (Figure 2g).

**Table 1.** Quantification of parthenogenesis and seed set in cowpea *pAtDD45::VuBBML1* and *pAtDD25::VuBBML1- pAtDD45::VuBBML1* transformants.

| Genotype | Number of plants analyzed | Average number of emasculated flowers per plant (total) | Average number of cytologically ovules analyzed (total) | Number of plants showing parthenogenesis in egg cell (EC) or central cell (CC) (%) |  | % Parthenogenesis range in egg cell (EC) or central cell (CC) (average) |  |
| --- | --- | --- | --- | --- | --- | --- | --- |
|  |  |  |  | EC | CC | EC | CC |
| *pAtDD45::VuBBML1* | 10 | 2.8 (28) | 34.4 (344) | 6 (60) | 0 | 0 – 15.8 (5.5) | 0 |
| *pAtDD25::VuBBML1-pAtDD45::VuBBML1* | 20 | 3.65 (73) | 41.6 (832) | 20 (100) | 11 (55) | 8.2 - 31.8 (19.8) | 0 – 9.4 (2.5) |

| Genotype | Number of plants analyzed | Average number of pods analyzed per plant (total) | Average number of harvested seeds per plant (%) |  | Average number of aborted seeds (%) | Average number of unfertilized ovules per plant |
| --- | --- | --- | --- | --- | --- | --- |
|  |  |  | Normal 2n | Small n |  |  |
| Untransformed | 2 | 5 (10) | 68.5 (99.2) | 0 | 1 (2) | 11 |
| *pAtDD45::VuBBML1* | 3 | 2.66 (8) | 14 (84) | 0 | 2.66 (16) | 25.33 |
| T0 *pAtDD25::VuBBML1-pAtDD45::VuBBML1* | 4 | 5.25 (21) | 45.5 (74.5) | 7.75 (12.7) | 7.75 (12.7) | 23 |
| T1 *pAtDD25::VuBBML1-pAtDD45::VuBBML1* | 6 | 4 (24) | 36 (66.4) | 10.16 (18.8) | 8 (14.8) | 9.83 |

**Table 2.** Transmission efficiency of the transgene in hemizygous *pAtDD25::VuBBML1-pAtDD45::VuBBML1* (*dBBML1*) plants.

|  | Total F1 progeny | dBML<sup>a</sup><br>(+) | dBML<sup>b</sup><br>(-) | dBML1 TE<sup>c</sup><br>(%) | DsRed<sup>d</sup><br>(-) | Viable parthenogenesis<br>(%) |
| --- | --- | --- | --- | --- | --- | --- |
| dBML1 (female) x DsRed (male) | 65 | 31 | 34 | 95.4 | 11 | 16.9 |
| DsRed (female) x dBML1 (male) | 53 | 25 | 28 | 94.3 | 0 | 0 |

<sup>a</sup> Transgenic *pAtDD25::VuBBML1-pAtDD45::VuBBML1* F1 plants
<sup>b</sup> Non-transgenic *pAtDD25::VuBBML1-pAtDD45::VuBBML1* F1 plants
<sup>c</sup> Transmission efficiency
<sup>d</sup> F1 progeny not showing *DsRed-Express* fluorescence

Collectively these results show that exclusive expression of *VuBBML1 or ToPAR* in the egg cell is not sufficient to induce viable haploid seed as parthenogenetic embryos arrest prior to the globular stage. Under self-pollinating conditions, free nuclear endosperm is not initiated. Following cross pollination, the transgene is only transmitted through male gametes, indicating that all parthenogenetic embryos arrest before reaching maturity in the absence of endosperm initiation. We propose that the exclusive activation of *VuBBML1* or *ToPAR* in the egg cell may impede fertilization of the central cell, preventing endosperm initiation and subsequently blocking progression of embryogenesis. The dual expression of *VuBBML1* in the egg cell and central cell might allow a synchronized crosstalk between the two gametophytic cell types. This crosstalk is likely necessary for simultaneously inducing both parthenogenesis and the first free nuclear divisions by either autonomous endosperm development or by allowing fertilization of the central cell. The elucidation of the mechanisms controlling endosperm development in developing seeds containing parthenogenetic embryos will require additional work.

In combination with meiotic avoidance, the recovery of viable haploid seeds through parthenogenesis is a necessary component for the establishment of synthetic apomixis in cowpea, a legume of major importance for sub-Saharan African countries. We expect that similar approaches will allow viable parthenogenesis in other legumes and eudicot species, opening the possibility for double haploid technologies to reduce the time and cost for obtaining fully homozygous inbred lines.

## Methods

### Plant material and growth conditions

Seeds of *Vigna unguiculata* IT86D-1010, Hyb18 and the *pGmEF1a::DsRed-Express* transgenic line were germinated and grown in 3.75-L pots filled with a soil mixture of 35% leaf coarse milled composted, 35% sunshine soil, 30% of perlite and fertilized with N-P-K:20-20-20. Plants were grown under greenhouse conditions, temperatures in spring-summer were 26–32°C and 22–27°C in autumn–winter with a relative humidity 50-70% during the day and 60-90% during night. For *in vitro* germination, seeds were surface sterilized using chlorine for 5 minutes, washed in sterile water, cultivated in bean germination medium (BGM) [24] for 24 hours in the dark, and transferred to MS medium supplemented with 1x MS vitamins and 30mg/L of meropenem. *In vitro* germination proceeded under controlled conditions of 16h/8h day/night photoperiod and 26°C. Plantlets were transferred directly to soil and grown under greenhouse conditions.

### Phylogenetic analysis of VuBBML1 family

Candidate *VuBBML1* genes were initially identified from early genomic sequences of IT86D10-10 and IT97K-499-35 [19] in addition to the IT97K-499-35 cowpea reference genome sequences [20, 22]. Genes with two APT2 motifs, BBM motifs an uANT2 motif an absence of an miR172 binding site, were sought given these features are characteristic of parthenogenesis inducing BBML/BBM genes from *Cenchrus squamulatus* (syn. *Pennisetum squamulatum*; PsASGR-BBML, EU559280), *Arabidopsis thaliana* (AtBBM, AT5G17430.1), *Oryza sativa* (OsBBM1, LOC_Os11g19060.1.p), and *Zea mays* (ZmBBM1, Zm00001eb247080_P001) were obtained from NCBI and Phytozome. Protein domains were predicted using NCBI Conserved Domain Search and Pfam. All APETALA2 (AP2) domain-containing proteins across the IT97K-499-35 cowpea reference genome were identified using HMMER v3.4 (http://hmmer.org/) with default settings. Protein sequence alignments were subsequently constructed using MAFFT v7.490 with the E-INS-I algorithm [25], employing the BLOSUM62 scoring matrix and a gap open penalty of 1.53. FastTree 2.1.11 [26] was used to construct a rooted maximum-likelihood tree with 3,000 rate categories of sites, including optimized Gamma20 likelihood and pseudocount settings. Statistical significance of branches was evaluated with the Shimodaira-Hasegawa (SH)-like local supports ratio test. The AP2 protein of cowpea that clustered with the verified BBM/BBML sequence VuBBMLVigun05g053500.1.p (see Figure S3A). The same approach was used to identify BBML homologues from other species, including *Medicago truncatula* (MtBBML1, Medtr7g080460.1.p), *Glycine max* (GmBBML1a, Glyma.18G244600.1.p and GmBBML1b, Glyma.09G248200.1.p), and *Sorghum bicolor* (SbBBML1, Sobic.004G214300.1.p). The final rooted phylogenetic tree was constructed by combining all cowpea double AP2 proteins with BBM proteins from the other seven species, along with the outgroup protein AtWUS (At2g17950.1). Visualization of the phylogenetic tree was generated using Geneious software and edited by Adobe Illustrator.

### Construction of plant transformation vectors

Gateway entry vectors containing cell-type specific promoter elements together with genes encoding fluorescent proteins *pAtRKD2:DsRed-Express*, *pAtDD45:DsRed-Express*, and *pAtDD25:DsRed-Express* were gifts from Shai Lawit and Marc Albertsen [27]. Construction of cowpea transformation vectors followed previously described procedures [24].

For construction of the *pToPAR::ToPAR* vector, a genomic sequence containing the promoter of *ToPAR* and the *ToPAR* gene was amplified from *Taraxacum officinale (*primers: ToPARF1 and ToParR; see Table S10). For construction of the *pDD45::VuBBML1* vector, the *pAtDD45* promoter was amplified from *Col-0 Arabidopsis thaliana* genomic DNA, and the *VuBBML1* gene was amplified from cDNA of *Vigna unguiculata (*primers: pAtDD45F and pAtDD45R; VuBBML1F and VuBBML1R; see Table S10). For construction of the *pAtRKD2::VuBBML1* (genomic) vector, an *ApaI* fragment consisting of 579 bp of the *AtRKD2 (*At1g74480) promoter, a 3800 bp genomic *VuBBML1* (Vu05g053500) fragment and 277 bp *NOS* terminator were placed into a modified RC677d vector [28] containing the soybean codon optimized *aadA1* gene for spectinomycin selection (UGAHyG010). The ApaI *pAtRKD2::VuBBML1* (genomic) fragment was separated from the *aadA1* cassette by a TBS insulator region. For construction of the *pRKD2::ToPAR* (cDNA) vector, the cDNA of *ToPAR* (544 bp) containing cloning enzyme restriction sites, was synthesized by Genescript Biotech Corporation (Piscataway, NJ, USA) and replaced the *VuBBML1* genomic fragment of UGAHyG010. This cassette was transferred into UQHyG017b, a vector containing a modified spectinomycin cassette for more efficient tissue culture selection (gift from Corteva) and a *pVuUB4::ZsGreen* cassette for visualization in transgenic plants and seedlings.

For the *pDD25::VuBBML-pDD45:VuBBML1* construct, the *pAtDD25* promoter was amplified from Arabidopsis *Col-0* genomic DNA (primers: pAtDD25F and pAtDD25R; see Table S6). The *VuBBML1* sequence was modified by selective amplification to remove internal BpiI and BsaI sites, introducing synonymous mutations on Gly-298 (GGA to GGG) and Ser-673 (TCT to TCC), respectively. All PCR fragments were cloned into the PCR8-TOPO vector and all recombination reactions were performed using Gateway™ LR Clonase™ II. A transcriptional unit containing a Gateway-compatible cassette fused to the *Arabidopsis thaliana* terminator tOCS was assembled by GoldenGate technology using BsaI sites as linkers. The backbone includes a plant selection spectinomycin cassette, the Gateway compatible cassette, and the backbone pAGM4723 from the MoClo kit as the final plasmid [29].Vectors were introduced into *Agrobacterium tumefaciens* LBA4404 Thy-strain, and transformation was performed to using immature embryo axes and regeneration protocols as previously described [24].

### Crossing and transgenic genotyping

For crosses, flowers at stage 1F-IX were emasculated and protected for 10h to 24h prior to pollination [19]. Pollination was conducted at early morning hours using freshly collected male flowers. For genotyping, DNA was extracted from leaves using the CTAB method [30], in some cases with minor modifications [28]. Leaves were frozen in liquid N_2_ and grinded to powder using Minibead beater (Biospec products) before addition of CTAB buffer and incubation at 55°C for 15 min. Following centrifugation, the supernatant was extracted with 24:1 chloroform: iso-amyl alcohol. before recovering the supernatant and adding 7.5 M ammonium acetate followed by absolute ethanol. DNA was collected recovered after centrifugation, washed with 70% ethanol, air-dried and dissolved in sterile DNase free water. DNA was treated with RNase A (Invitrogen). Genotyping analysis was carried out with primers shown in Table S6 using the following PCR cycle parameters: 94°C for 30 s, 60°C for 30 s 72°C for 1 min for 35 cycles.

### Whole genome sequencing and genotyping

To recover high molecular weight DNA, lysate was extracted from cowpea plants using the modified CTAB method. Lysate was added with absolute ethanol (1/1) and it was placed into the DNA spin column (QIAGEN) and processed as indicated by the supplier (DNeasy Blood & tissue kit, QIAGEN, CA). Quality parameters were assessed with nanodrop (Invitrogen). Sequencing reads were produced using the IIlumina NovaSeq platform. After quality filtering, reads were mapped to the publicly available *Vigna unguiculata* genome [20] and single nucleotide polymorphisms (SNPs) identified in regions with at least 10X depth coverage, using *SAMtools.* Heterozygous SNPs were mapped, and the chromosomal position was identified in maternal genomes prior to comparison to haploid progeny candidates at all eleven chromosomes.

### Ploidy measurement

Approximately 1 cm^2^ of fresh leaf tissue was chopped with 2 ml of lysis buffer. The mixture was filtered through a 30-µm nylon filter. Nuclei were stained with propidium iodide for 20 min in the dark (lysis buffer and propidium iodide from the Cystain-PI Oxprotect kit). A PARTEC CyFlow-Cube6 flow cytometer was used to evaluate DNA content. Data analysis, including gating was conducted using ModFit LT v3.3 software.

### Cytological analysis

Manual emasculation was performed using tweezers to remove anthers and stigma from flowers at 1F-IX stage, before anthesis and pollen dehiscence. Three days after emasculation, individual gynoecia were dissected under a stereoscope to expose the ovules without detaching them from the placenta and fixed in FAA (50 % ethanol, 10% formaldehyde, 4% acetic acid and water) for at least 12 h. Strings of ovules were washed in 70% ethanol. Light microscopy was performed by clearing strings of ovules with Herr’s solution and incubated for 24h at RT. Ovules were mounted in the same clearing solution and observed under Nomarski illumination using a DMR Leica microscope. Confocal microscopy was performed by clearing through ethanol dehydration and methyl salicylate [31]. Cleared ovules were mounted in methyl salicylate and observed with a Nikon C2+ confocal microscope using a solid state green (488 nm laser line/emission 400-480) or red (561 nm laser line/emission 544-624) laser/filter. For reporter gene expression of *pAtDD25* and *pAtDD45* activity, dissected ovules were mounted in water and observed with a SP5 Leica confocal microscope and with a 558 nm laser line for excitation of the *DsRed-Express* fluorescent protein; emission was at 565-720 nm. For excitation of SR2200, emission was at 405 nm and detection at 410-480 nm. In the case of d*BBML1* x *pGmEF1a::DsRed-Express* crosses, fluorescence analysis of F1 progeny was performed in embryos after removing the seed coat and analyzed under a stereomicroscope equipped with the NIGHTSEA fluorescence adapter and a Green filter (510-540 nm) from Electron Microscopy Sciences. For chromosome observation, we adapted a previously described root tip squash method [32]. *AmCyan1* signals were observed with Zeiss filter set 46; excitation was at 468 nm and emission at 489 nm. Root meristem cells were collected from actively growing roots of *in vitro* germinated seedlings or young plants and pre-treated with 2 mM 8-hydroxyquinoline for 3 h at RT; then transferred to a fixative Farmer’s solution (3:1 v/v ethanol:glacial acetic acid) and incubated for 24 h at RT. Samples were digested in an enzyme mixture for 60 min at 37 °C. The steam drop method was used to prepare chromosome spreads [32]. Samples were stained with 4′,6-diamidinophenylindole (DAPI), mounted on a slide in Vectashield® and observed under a Leica SPE confocal laser scanning microscope using settings for Alexa 488 at 60X.

## Availability of materials

Biological materials generated in the course of this work are available from the authors upon reasonable request.

## Data availability

The data supporting the findings of this work are available within the paper, its extended data Figures and Tables included in the Supplementary Information. Sequences from Arabidopsis and cowpea are available through TAIR (https://www.arabidopsis.org/) and Phytozome (https://phytozome-next.jgi.doe.gov/). Accession numbers are also provided in the relevant sections of the paper. Source data are provided with this paper.

## Acknowledgments

We thank Shai Lawit and Marc Albertsen for the generous gift of Arabidopsis promoter fluorescent marker fusion vectors; members of the CSIRO Capturing Heterosis team, Tracy How, Dilrukshi Nagahatenna and Natalia Bazanova for technical support; and Mireya Hernandez, Ana Lilia Solorzano and Angie Strelow for administrative support.

## Contributions

IAM, ORM, GLM and JPVC designed and performed experiments giving rise to viable pathenogenesis; JC, SB, QOV, HS, SaBh, RH, NG and JdC provided supporting information and peformed companion and confirmation experiments; JC, KR, JL and StBe performed plant tranformation experiments; MR, MJ, MH, and SDJ indentified cowpea genes and promoters; MVE performed computational analysis; BF, UG and POA supervised and analyzed experiments; UG, POA, BF, AMGK and JPVC conceived the project; AMGK and JPVC wrote the paper with input from other authors.

## Supplementary Information

**Figure S1.**
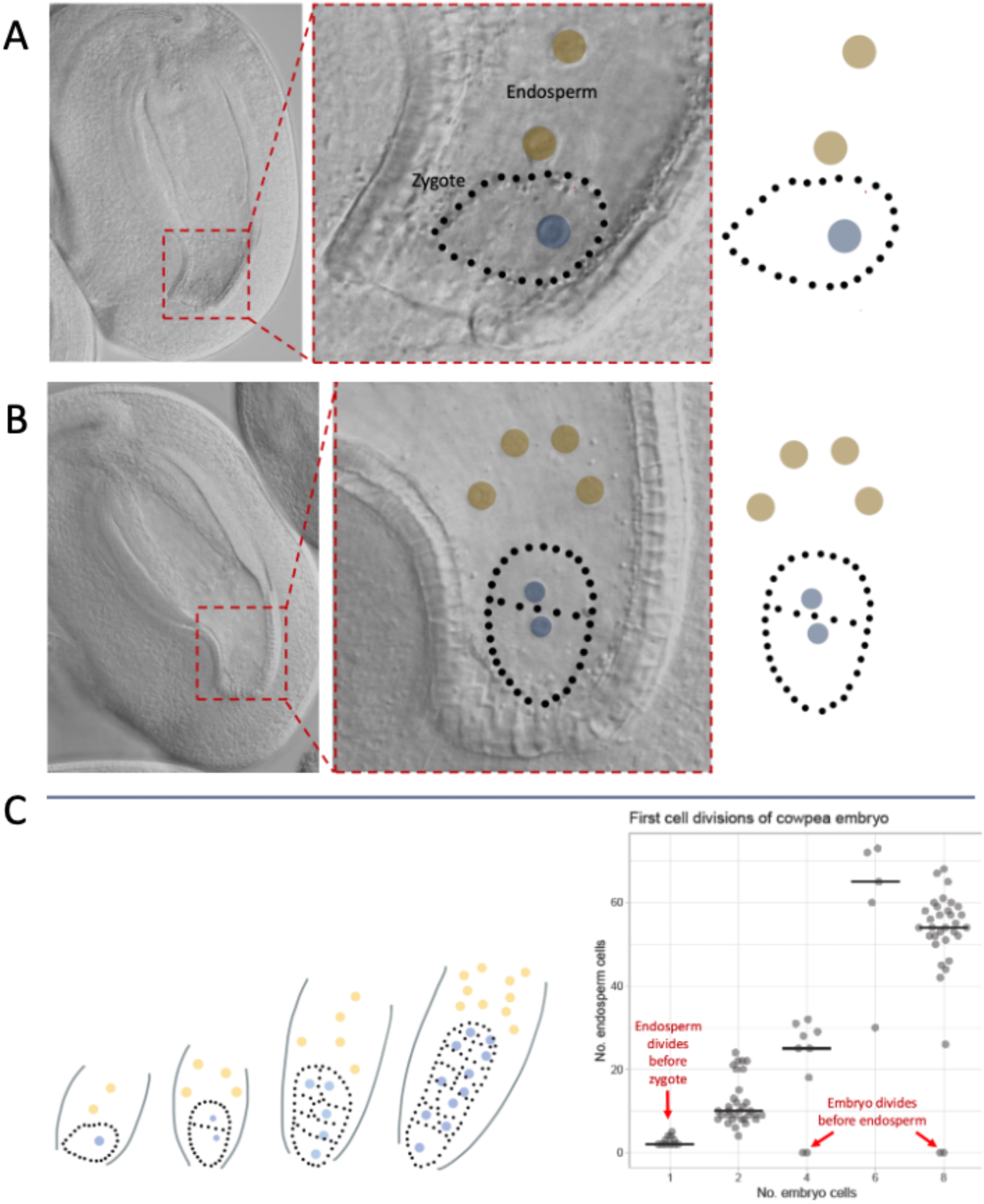
Examples of coordinated divisions of the zygote and primary endosperm nucleus following double fertilization in cowpea. (**A**) Female gametophyte at 10 hours after pollination showing an undivided zygote and the first division of the primary endosperm nucleus. (**B**) Female gametophyte at 10 hours after pollination showing a 2-cell embryo and four endosperm nuclei. (**C**) Kinetic model showing coordinated progression of embryo and endosperm development based on 97 embryo sacs observed 10 to 24 hours after pollination.

**Figure S2.**
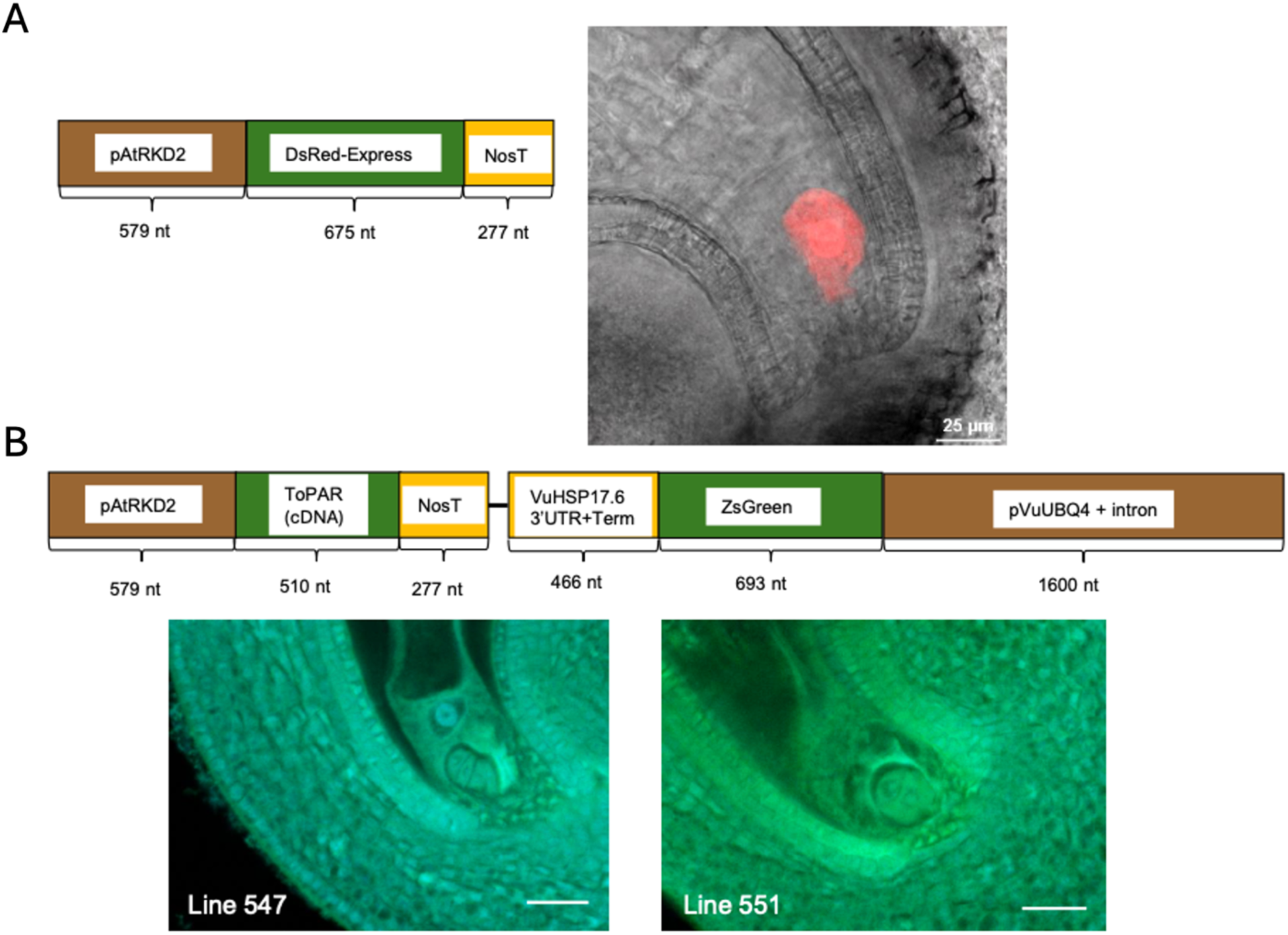
**Parthenogenesis in *pAtRKD2::ToPAR* transformants**. (**A**) Transgene cassette and expression of *pAtRKD2::DsRed-Express* within the egg cell of a cowpea embryo sac fixed from an open flower prior to fertilization. (**B**) Transgene cassette and phenotype of parthenogenic embryo development from ovules fixed from emasculated mature, open flowers containing the *pAtRKD2::ToPAR* construct.

**Figure S3.**
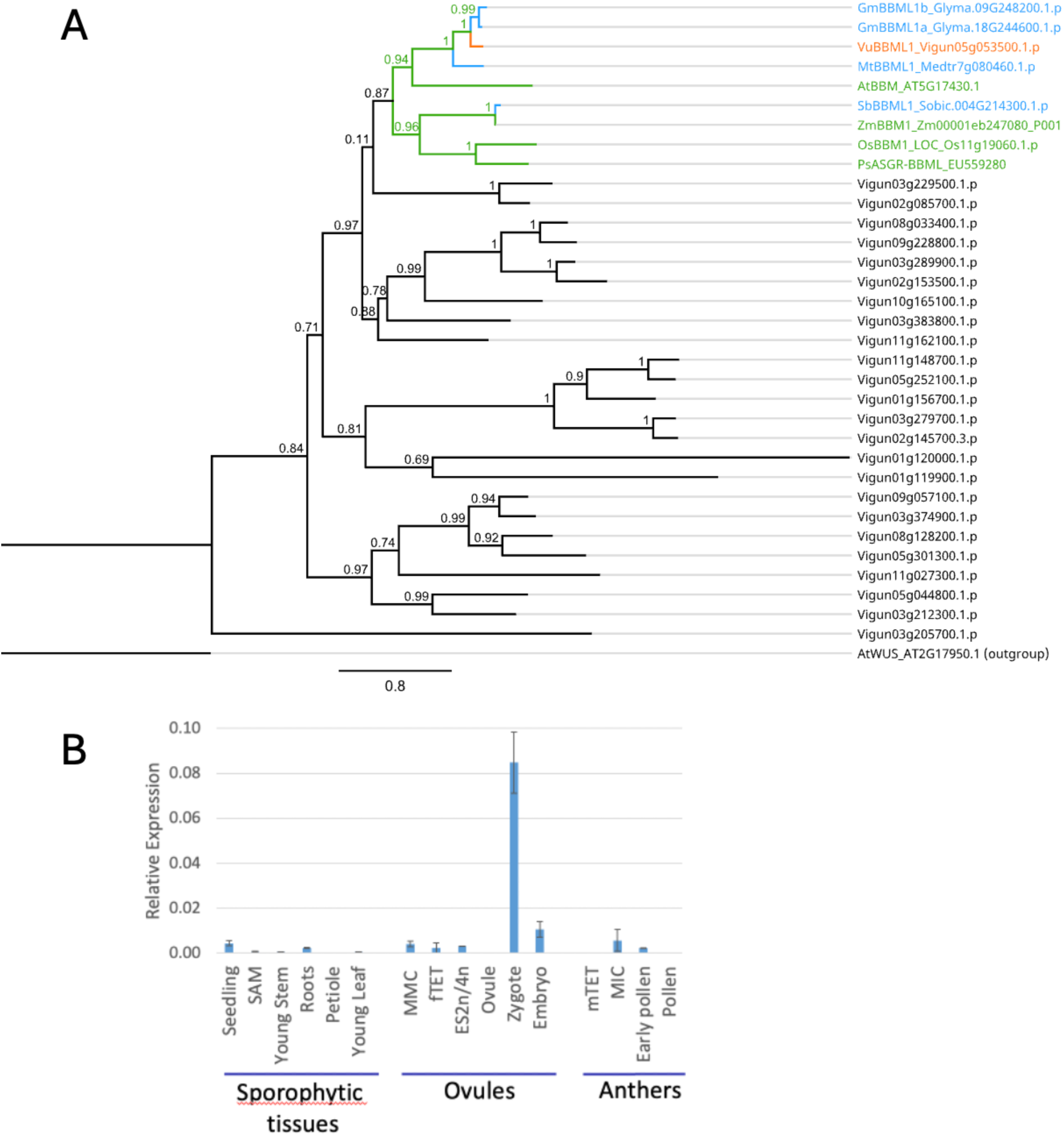
Phylogenetic tree of cowpea double AP2 domain proteins and pattern of expression of *VuBBML1* in vegetative and reproductive tissues. (A) A Maximum Likelihood (ML) tree was constructed using amino acid sequences of 25 double AP2 domains in cowpea, along with BBM proteins from other species, including soybean (*Glycine max*), *Medicago truncatula*, *Arabidopsis thaliana*, sorghum (*Sorghum bicolor*), rice (*Oryza sativa*), and *Cenchrus squamulatum* (syn. *Pennisetum squamulatum*). The analysis was conducted using FastTree version 2.1.11, with AtWUS protein serving as an outgroup. Shimodaira-Hasegawa (SH)-like local support values are indicated at each branch. The VuBBML1 protein is highlighted in orange, while other previously reported BBM-like proteins are marked in green; newly predicted BBM-like proteins from additional legume species and *Sorghum bicolor* are marked in blue; additional AP2 domain proteins from cowpea are marked in black. (B) qRT-PCR of *VuBBML1* in different cowpea tissues, including vegetative and reproductive organs. MMC= megaspore mother cell; SAM = shoot apical meristem; fTET = functional megaspore stage; ES2n/4n = 2-nuclear to 4-nuclear female gametophytic stage; mTET = male tetrad stage; MIC= microsporocyte stage.

**Figure S4.**
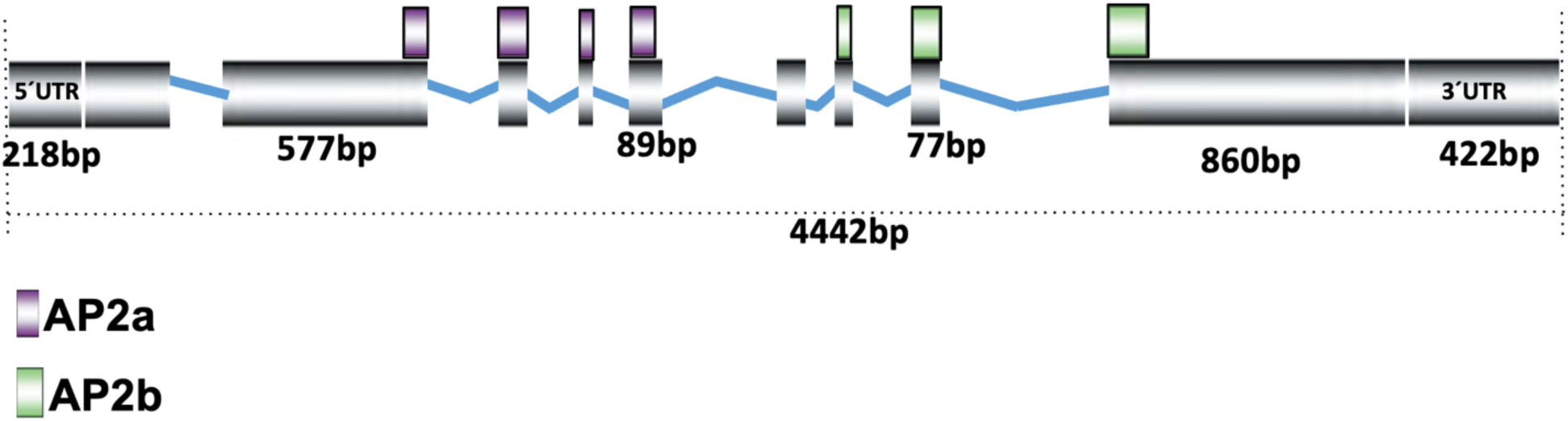
Genomic structure of the *Vigna unguiculata BABYBOOM-LIKE1* gene (*VuBBML1*; Vu05g053500). Exons are represented by gray rectangles and introns by blue lines. The distribution of two AP2 domain coding regions is indicated above the corresponding exons.

**Figure S5.**
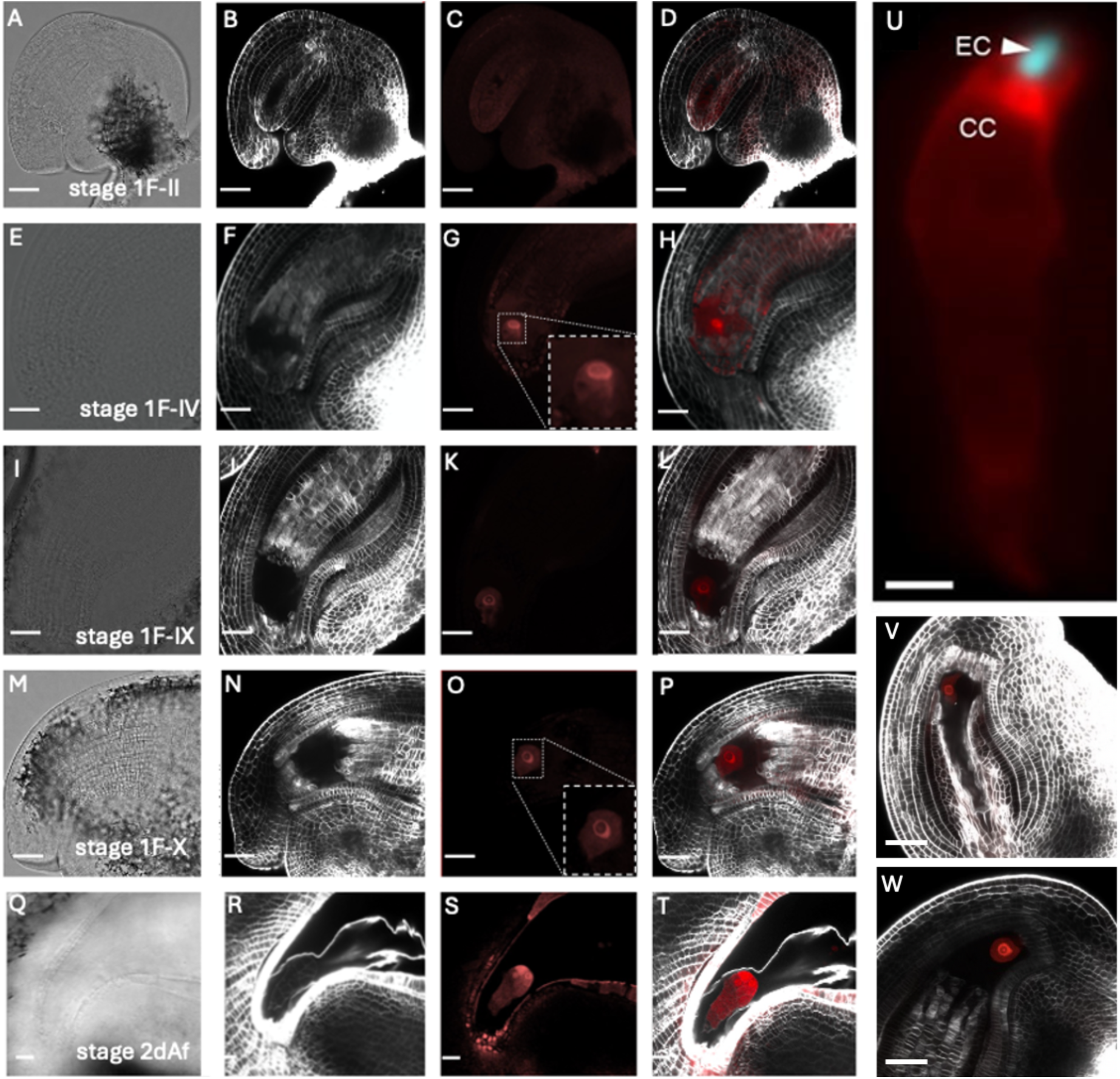
Specific activity of the Arabidopsis *pAtRKD2*, *pAtDD45* and *pAtDD25* promoters in developing ovules of cowpea (see also ref. 20). Cowpea ovules of plants expressing the construct *AtDD45::DsRed-Express* at different stages of development. Stages according to [18] are indicated in the bottom-right corner of the panels in the first column. For each stage, bright field (A,E,I,M,Q), SR2200 staining of the cell wall (B,F,L,N,R), *DsRed-Express* fluorescence (C,G,K,O,S), and a combination of the latter two are shown (D,H,L,P,T). Note that there is some autofluorescence in the sporophytic cells of the ovule. Detection of the *DsRed-Express* signal in the egg cell starts at stage 1F-IV (G) and is still detectable in the embryo at 2 days after fertilization (daf) (S). In G and O, the signal in the egg and the zygote, respectively, has been enlarged and is shown in the right corner of the panel (dotted lines). (U) *pAtDD25:DsRed-Express* (red) in the fertilized central cell and *pAtEC1.1:AmCyan1* (cyan) in fertilized egg cell. (V) *pAtRKD2:DsRed-Express* in the unfertilized egg cell at stage 1F-VII. (W) *pAtRKD2:DsRed-Express* (red) in the unfertilized egg cell at stage 1F-X. Scale bars in A to T, V and W: 10 µm. Scale bar in U: 50 µm.

**Figure S6.**
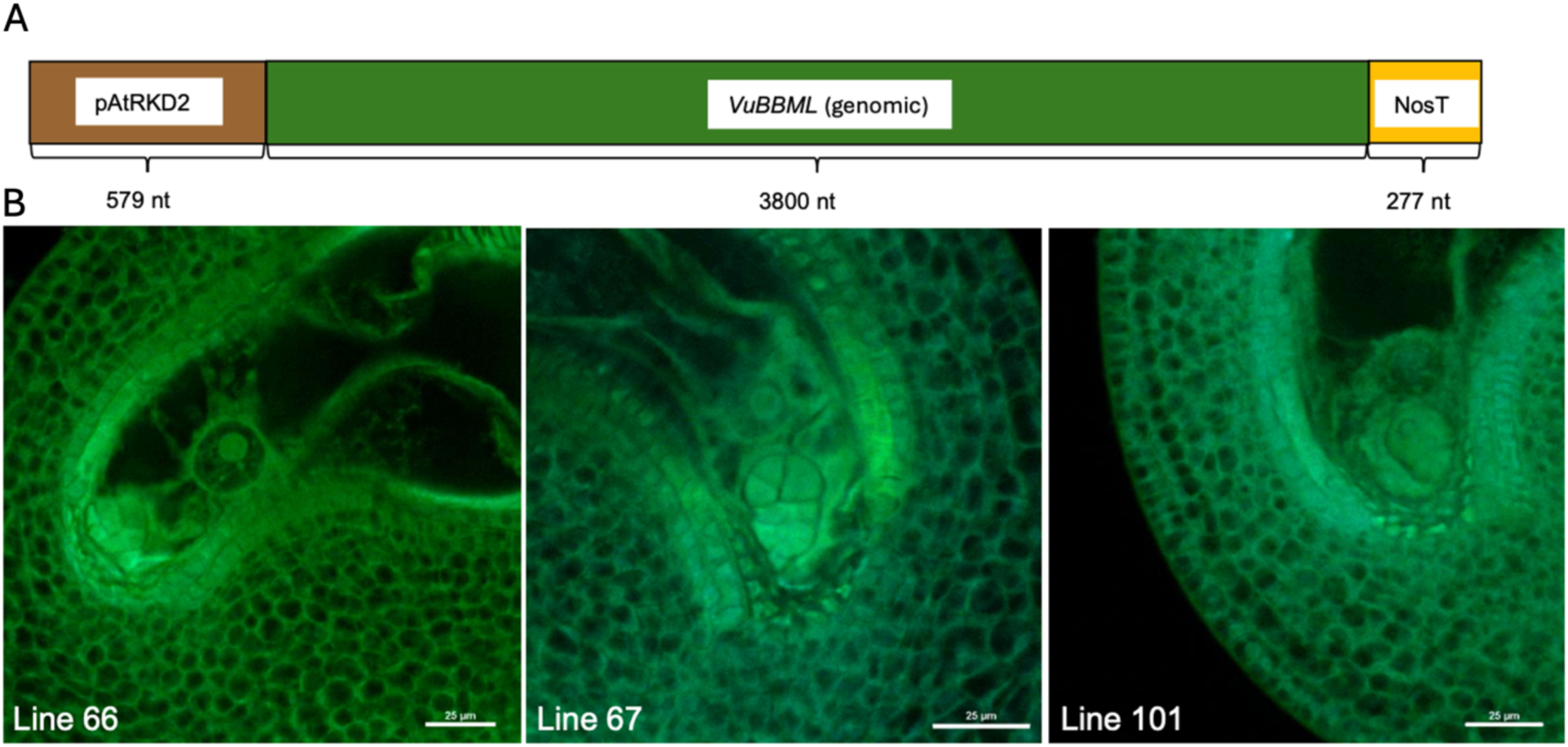
Parthenogenesis in *pAtRKD2::VuBBML1* transformants. (A) Transgene construct combining the Arabidopsis pAtRKD2 promoter and the VuBBML1 gene from cowpea. (**B**) parthenogenic embryo development in ovules from emasculated mature flowers of *pAtRKD2::VuBBML1* transformants. Micrographs show parthenogenetic embryos in ovules from three different transformant lines.

**Figure S7.**
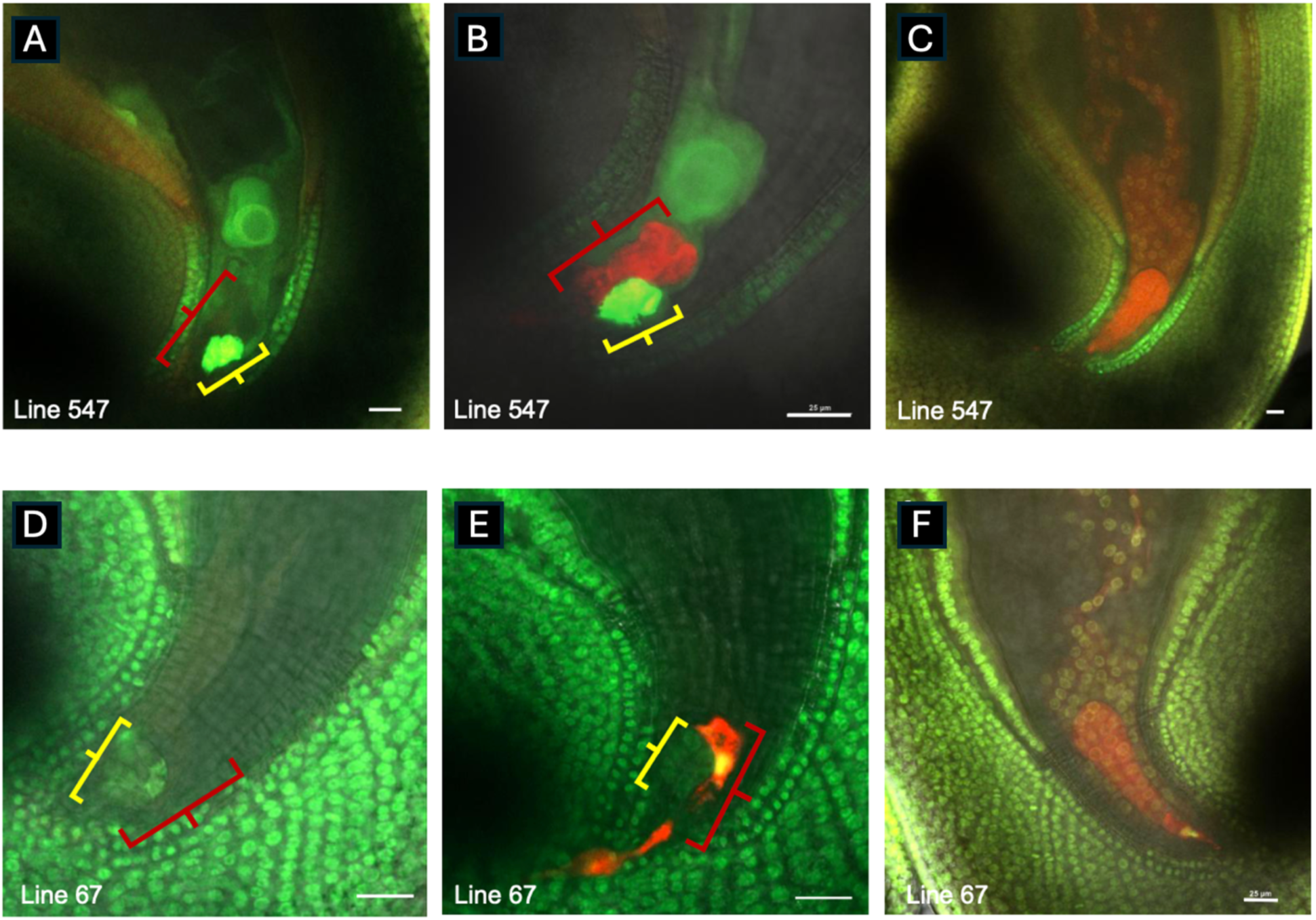
Pollen tube attraction in the ovule of *pAtRKD2::ToPAR* and *pAtRKD2::VuBBML1* transformants. Three outcomes result for the pollination of p*AtRKD2::ToPAR* (cDNA) T_2_ line 547 (**A** to **C**) and *pAtRKD2::VuBBML1* (genomic) T_2_ line 67 (**D** to **F**) with homozygous *DsRed-Express* pollen. Both maternal lines are heterozygous for the transgene. Green fluorescence in *pAtRKD2::ToPAR* (cDNA) T_2_ line 547 is due to expression of a *pVuUB4::ZsGreen* cassette (shown in figures A to C) included in the *pAtRKD2::ToPAR (cDNA)* vector; while in *RKD2p::VuBBML1* (genomic) line 67, green fluorescence is caused by ovule staining with a pico-green DNA dye (shown in figures D to F). The pico-green DNA dye helps visualize the parthenogenetic embryos, is not transgene specific, and is present all cells to varying degrees. (**A and D**) Ovules containing parthenogenetic embryos (yellow bracket) but without a *DsRed-Express* signal from a pollen tube within the synergid (red bracket). (**B and E**) Ovules containing a parthenogenetic embryo with a *DsRed-Express* signal from a pollen tube within the synergid. (**C and F**) Ovules without the *pAtRKD2::ToPAR* (cDNA) or *pAtRKD2p::VuBBML1* (genomic) transgene displaying wild-type embryo and endosperm development. Scale bar: 25µm.

---

**Table S1.** Number of cowpea embryo cells and free nuclei in the endosperm 10 to 24 hours after pollination.

| Sample No. | Number of Embryo Cells | Number of Free Nuclei in Endosperm |
| --- | --- | --- |
| 10h_sample_16 | 1 | 5 |
| 10h_sample_29 | 1 | 2 |
| 10h_sample_3 | 1 | 4 |
| 10h_sample_7 | 1 | 2 |
| 10h_sample_8 | 1 | 4 |
| 13h_sample_10 | 1 | 2 |
| 13h_sample_11 | 1 | 2 |
| 13h_sample_14 | 1 | 2 |
| 13h_sample_16 | 1 | 2 |
| 13h_sample_4 | 1 | 3 |
| 13h_sample_5 | 1 | 3 |
| 10h_sample_1 | 2 | 10 |
| 10h_sample_10 | 2 | 7 |
| 10h_sample_11 | 2 | 8 |
| 10h_sample_12 | 2 | 9 |
| 10h_sample_13 | 2 | 11 |
| 10h_sample_14 | 2 | 10 |
| 10h_sample_15 | 2 | 12 |
| 10h_sample_17 | 2 | 10 |
| 10h_sample_18 | 2 | 9 |
| 10h_sample_19 | 2 | 10 |
| 10h_sample_2 | 2 | 12 |
| 10h_sample_20 | 2 | 11 |
| 10h_sample_21 | 2 | 15 |
| 10h_sample_22 | 2 | 9 |
| 10h_sample_23 | 2 | 7 |
| 10h_sample_24 | 2 | 8 |
| 10h_sample_25 | 2 | 8 |
| 10h_sample_26 | 2 | 8 |
| 10h_sample_27 | 2 | 22 |
| 10h_sample_28 | 2 | 9 |
| 10h_sample_4 | 2 | 8 |
| 10h_sample_5 | 2 | 9 |
| 10h_sample_6 | 2 | 4 |
| 10h_sample_9 | 2 | 13 |
| 13h_sample_12 | 2 | 24 |
| 13h_sample_13 | 2 | 9 |
| 13h_sample_15 | 2 | 20 |
| 13h_sample_3 | 2 | 6 |
| 13h_sample_6 | 2 | 20 |
| 13h_sample_7 | 2 | 22 |
| 13h_sample_8 | 2 | 21 |
| 13h_sample_9 | 2 | 22 |
| 13h_sample_1 | 4 | 18 |
| 13h_sample_2 | 4 | 25 |
| 24h_sample_17 | 4 | 0 |
| 24h_sample_18 | 4 | 0 |
| 24h_sample_19 | 4 | 29 |
| 24h_sample_22 | 4 | 32 |
| 24h_sample_23 | 4 | 31 |
| 24h_sample_50 | 4 | 25 |
| 24h_sample_9 | 4 | 28 |
| 24h_sample_12 | 6 | 65 |
| 24h_sample_15 | 6 | 60 |
| 24h_sample_16 | 6 | 72 |
| 24h_sample_2 | 6 | 73 |
| 24h_sample_26 | 6 | 30 |
| 24h_sample_1 | 8 | 53 |
| 24h_sample_11 | 8 | 42 |
| 24h_sample_13 | 8 | 58 |
| 24h_sample_14 | 8 | 54 |
| 24h_sample_20 | 8 | 61 |
| 24h_sample_21 | 8 | 60 |
| 24h_sample_24 | 8 | 0 |
| 24h_sample_25 | 8 | 52 |
| 24h_sample_27 | 8 | 58 |
| 24h_sample_28 | 8 | 26 |
| 24h_sample_29 | 8 | 55 |
| 24h_sample_30 | 8 | 44 |
| 24h_sample_31 | 8 | 54 |
| 24h_sample_32 | 8 | 45 |
| 24h_sample_33 | 8 | 0 |
| 24h_sample_34 | 8 | 56 |
| 24h_sample_35 | 8 | 57 |
| 24h_sample_37 | 8 | 59 |
| 24h_sample_39 | 8 | 60 |
| 24h_sample_41 | 8 | 54 |
| 24h_sample_42 | 8 | 67 |
| 24h_sample_43 | 8 | 57 |
| 24h_sample_44 | 8 | 51 |
| 24h_sample_45 | 8 | 54 |
| 24h_sample_46 | 8 | 52 |
| 24h_sample_47 | 8 | 59 |
| 24h_sample_48 | 8 | 52 |
| 24h_sample_49 | 8 | 50 |
| 24h_sample_5 | 8 | 68 |
| 24h_sample_51 | 8 | 46 |
| 24h_sample_52 | 8 | 57 |
| 24h_sample_6 | 8 | 53 |
| 24h_sample_8 | 8 | 65 |
| 24h_sample_10 | 10 | 62 |
| 24h_sample_7 | 10 | 66 |
| 24h_sample_40 | 11 | 55 |
| 24h_sample_3 | 12 | 55 |
| 24h_sample_36 | 14 | 73 |
| 24h_sample_38 | 14 | 76 |
| 24h_sample_4 | 14 | 62 |
| 12h sample L1 | 1 | 6 |
| 12h sample L2 | 1 | 4 |
| 12h sample L3 | 1 | 4 |
| 12h sample L4 | 1 | 2 |
| 12h sample L5 | 1 | 5 |
| 12h sample L6 | 1 | 6 |
| 124h sample L7 | 1 | 6 |
| 12h sample L8 | 1 | 4 |
| 12h sample L9 | 1 | 2 |
| 12h sample L10 | 1 | 0 |
| 12h sample L11 | 1 | 4 |
| 12h sample L12 | 1 | 4 |
| 12h sample L13 | 1 | 6 |
| 12h sample L14 | 1 | 6 |
| 12h sample L15 | 1 | 4 |
| 12h sample L16 | 1 | 2 |
| 12h sample L17 | 1 | 2 |
| 12h sample L18 | 1 | 4 |
| 12h sample L19 | 1 | 4 |
| 12h sample L20 | 1 | 6 |
| 12h sample L21 | 2 | 6 |
| 12h sample L22 | 2 | 10 |
| 12h sample L23 | 2 | 0 |
| 12h sample L24 | 2 | 16 |
| 12h sample L25 | 2 | 8 |
| 12h sample L26 | 2 | 12 |
| 12h sample L27 | 2 | 10 |
| 12h sample L28 | 2 | 8 |
| 12h sample L29 | 2 | 17 |
| 12h sample L30 | 2 | 8 |
| 12h sample L31 | 2 | 10 |
| 12h sample L32 | 2 | 12 |
| 12h sample L33 | 2 | 15 |
| 12h sample L34 | 2 | 10 |
| 12h sample L35 | 2 | 0 |
| 12h sample L36 | 2 | 8 |
| 12h sample L37 | 4 | 11 |
| 12h sample L38 | 4 | 10 |
| 12h sample L39 | 4 | 16 |
| 12h sample L40 | 4 | 22 |
| 12h sample L41 | 4 | 13 |
| 12h sample L42 | 4 | 21 |
| 12h sample L43 | 4 | 10 |
| 12h sample L44 | 4 | 12 |
| 12h sample L45 | 4 | 8 |
| 12h sample L46 | 4 | 11 |
| 24h sample L47 | 4 | 23 |
| 24h sample L48 | 4 | 26 |
| 24h sample L49 | 4 | 17 |
| 24h sample L50 | 4 | 14 |
| 24h sample L51 | 4 | 10 |
| 24h sample L52 | 4 | 17 |
| 24h sample L53 | 4 | 21 |
| 24h sample L54 | 8 | 36 |
| 24h sample L55 | 8 | 41 |
| 24h sample L56 | 8 | 25 |
| 24h sample L57 | 8 | 29 |
| 24h sample L58 | 8 | 22 |
| 24h sample L59 | 8 | 48 |
| 24h sample L60 | 8 | >50 |
| 24h sample L61 | 8 | >50 |
| 24h sample L62 | 8 | >50 |
| 24h sample L63 | 8 | >50 |
| 24h sample L64 | 8 | >50 |
| 24h sample L65 | 8 | >50 |
| 24h sample L66 | 8 | >50 |

**Table S2.** Analysis of parthenogenesis in *pToPAR::ToPAR* transformants.

| Plant ID | Number of emasculated flowers | Number of ovules analyzed | Number of ovules showing parthenogenesis (%) |
| --- | --- | --- | --- |
| PAR397 | 4 | 48 | 0 |
| PAR398 | 3 | 40 | 0 |
| PAR406 | 3 | 41 | 0 |
| PAR407 | 3 | 42 | 0 |
| PAR408 | 8 | 80 | 1 (1.25) |
| PAR409 | 10 | 97 | 0 |
| PAR415 | 4 | 56 | 2 (3.57) |
| PAR416 | 7 | 98 | 0 |
| PAR417 | 3 | 36 | 0 |
| PAR418 | 2 | 25 | 0 |
| PAR419 | 4 | 51 | 0 |
| PAR420 | 3 | 34 | 0 |
| PAR421 | 4 | 46 | 0 |
| PAR425 | 5 | 49 | 0 |
| PAR427 | 7 | 82 | 3 (3.65) |
| PAR428 | 6 | 74 | 0 |
| PAR429 | 7 | 90 | 3 (3.33) |
| PAR432 | 2 | 28 | 0 |
| PAR433 | 4 | 40 | 0 |
| PAR417 | 8 | 48 | 0 |
| PAR418 | 10 | 40 | 0 |
| PAR419 | 4 | 41 | 0 |
| PAR420 | 7 | 42 | 0 |
| PAR421 | 3 | 80 | 1 (1.25) |
| **Total** | 121 | 1308 | 10 (0.76) |

**Table S3.** Absence of p*AtRKD2::ToPAR (cDNA)* maternal inheritance among sexually derived diploid offspring from cross pollination with homozygous *pGmEF1A::DsRed-Express* pollen.

| Lines for crossing | Generation of maternal plant | # of plantlets tested | DsRed positive | ToPar/Spec positive |
| --- | --- | --- | --- | --- |
| 549-1 | T1 | 10 | 10 | 0 |
| 549-2 | T1 | 22 | 22 | 0 |
| Total |  | 32 | 32 | 0 |
| 547-1-2 | T2 | 4 | 4 | 0 |
| 547-1-3 | T2 | 34 | 34 | 0 |
| 547-1-4 | T2 | 5 | 5 | 0 |
| 547-2-1 | T2 | 6 | 6 | 0 |
| 547-2-3 | T2 | 5 | 5 | 0 |
| Total |  | 54 | 54 | 0 |

**Table S4.** Parthenogenesis efficiency and embryo developmental developmental stages in ovules of hybrid plants derived from crosses of a *pAtRKD2::VuBBML1* transformant (line 66) with different cowpea varieties. Only one (*) out of 38 F_1_ individuals showed a statistically significant increase in parthenogenesis (P<0.05) compared with the maternal transgenic line; as tested with a binomial generalized linear model (GLM) that estimated the log-odds of parthenogenetic ovule formation for each cross.

| Cross | F1 plant ID | Total ovules screened | Number of ovules with parthenogenesis (PE) | % PE | Number of embryonic cells | Collapsed or abnormal female gametophytes |
| --- | --- | --- | --- | --- | --- | --- |
| Bambey23x66 | 1 | 44 | 28 | 63% | 2-8 | 1 |
|  | 8 | 55 | 27 | 49% | 4-8 | 0 |
|  | 12 | 46 | 16 | 34% | 4-6 | 3 |
|  | 18 | 41 | 20 | 48% | 3-8 | 1 |
|  | 19 | 50 | 26 | 52% | 4-8 | 1 |
|  | 20 | 52 | 27 | 52% | 4-8 | 1 |
|  | 23 | 44 | 22 | 50% | 4-7 | 2 |
| PI 583259 x66 | 2 | 25 | 15 | 62% | 3-5 | 1 |
|  | 6 | 43 | 24 | 55% | 2-6 | 0 |
|  | 14 | 30 | 11 | 37% | 3-6 | 2 |
|  | 17 | 52 | 18 | 35% | 4-7 | 0 |
| TVu249x66 | 1 | 30 | 10 | 33% | 4-5 | 0 |
|  | 2 | 34 | 13 | 38% | 3-5 | 0 |
|  | 5* | 41 | 30 | 73% | 4-7 | 0 |
|  | 6 | 47 | 23 | 49% | 4-5 | 0 |
|  | 8 | 42 | 20 | 45% | 4-8 | 0 |
|  | 9 | 45 | 23 | 51% | 4-10 | 0 |
|  | 10 | 54 | 24 | 44% | 4-10 | 2 |
|  | 12 | 54 | 27 | 50% | 4-12 | 1 |
|  | 13 | 51 | 23 | 45% | 2-10 | 2 |
|  | 14 | 57 | 28 | 49% | 4-10 | 1 |
|  | 16 | 55 | 25 | 44% | 4-12 | 0 |
|  | 18 | 55 | 19 | 37% | 4-12 | 2 |
|  | 19 | 51 | 24 | 47% | 6-12 | 0 |
|  | 20 | 59 | 28 | 47% | 5-10 | 2 |
|  | 22 | 50 | 25 | 50% | 6-10 | 2 |
|  | 24 | 55 | 27 | 49% | 5-12 | 1 |
|  | 25 | 58 | 34 | 58% | 4-10 | 1 |
| TVu1503x66 | 2 | 58 | 29 | 50% | 2-10 | 1 |
|  | 3 | 52 | 28 | 53% | 4-10 | 0 |
|  | 4 | 44 | 19 | 43% | 4-8 | 7 |
|  | 6 | 53 | 30 | 56% | 4-10 | 2 |
|  | 8 | 46 | 16 | 35% | 2-10 | 1 |
|  | 11 | 58 | 32 | 55% | 2-10 | 5 |
|  | 13 | 44 | 24 | 54% | 2-14 |  |
|  | 20 | 61 | 35 | 57% | 2-10 | 3 |
|  | 23 | 46 | 22 | 52% | 2-8 | 0 |
|  | 24 | 56 | 30 | 53% | 2-10 | 2 |
| CW66 | T4-Plant<br>3 | 40 | 18 | 45% | 4-10 | 4 |
|  | T4-Plant<br>7 | 50 | 22 | 46% | 4-8 | 2 |
|  | T4-plant<br>10 | 40 | 19 | 47% | 4-10 | 4 |

**Table S5.** Quantification of parthenogenesis in ovules of T0 plants three days after emasculation.

| Plant Id | Number of emasculated flowers | Number of ovules cytologically analyzed | Number of ovules showing parthenogenesis in the egg cell (%) |
| --- | --- | --- | --- |
| sBBML1-764 | 4 | 49 | 0 |
| sBBML1-766 | 2 | 28 | 0 |
| sBBML1-775 | 2 | 24 | 1 (4.2) |
| sBBML1-777 | 4 | 42 | 0 |
| sBBML1-778 | 3 | 34 | 4 (11.8) |
| sBBML1-781 | 3 | 38 | 6 (15.8) |
| sBBML1-787 | 2 | 30 | 3 (10) |
| sBBML1-788 | 2 | 23 | 2 (8.7) |
| sBBML1-802 | 3 | 36 | 3 (8.3) |
| sBBML1-815 | 3 | 40 | 0 |
| **Total** | **28** | **344** | **19 (5.5)** |

*T0 pAtDD25::VuBBML1-pAtDD45::VuBBML1*
| Plant ID | Number of emasculated flowers | Number of ovules analyzed | Ovules showing parthenogenesis in EC (%) | Ovules showing parthenogenesis in CC (%) |
| --- | --- | --- | --- | --- |
| dBBML1-690 | 7 | 88 | 25 (28.4) | 6 (6.8) |
| dBBML1-699 | 5 | 48 | 11 (22.9) | 1 (2.1) |
| dBBML1-700 | 4 | 42 | 9 (21.4) | 1 (2.4) |
| dBBML1-708 | 6 | 61 | 12 (19.7) | 0 |
| dBBML1-709 | 4 | 42 | 5 (11.9) | 1 (2.4) |
| dBBML1-713 | 3 | 32 | 3 (9.4) | 3 (9.4) |
| dBBML1-791 | 3 | 33 | 10 (30.3) | 0 |
| dBBML1-792 | 4 | 46 | 5 (10.9) | 1 (2.2) |
| dBBML1-798 | 5 | 45 | 9 (20) | 0 |
| dBBML1-800 | 2 | 22 | 7 (31.8) | 0 |
| dBBML1-801 | 2 | 28 | 4 (14.3) | 1 (3.6) |
| dBBML1-816 | 5 | 59 | 14 (23.7) | 0 |
| dBBML1-817 | 2 | 26 | 3 (11.5) | 1 (3.8) |
| dBBML1-819 | 4 | 49 | 4 (8.2) | 0 |
| dBBML1-820 | 4 | 41 | 8 (19.5) | 0 |
| dBBML1-821 | 3 | 38 | 7(18.4) | 2 (5.2) |
| dBBML1-822 | 2 | 22 | 3 (13.6) | 0 |
| dBBML1-824 | 2 | 29 | 9 (31.1) | 2 (6.9) |
| dBBML1-831 | 3 | 38 | 8(21.1) | 0 |
| dBBML1-832 | 3 | 43 | 9(20.9) | 2 (4.6) |
| **Total** | **73** | **832** | **165 (19.8)** | **21 (2.5)** |

**Table S6.** Absence of *pAtRKD2::VuBBML1* maternal inheritance among sexually derived diploid offspring from cross pollination with homozygous *pGmEF1A::DsRed-Express* pollen.

| Lines for crossing | Generation of maternal plant | Number pf plantlets tested | DsRed positive | ToPar/Spec positive |
| --- | --- | --- | --- | --- |
| 66-9 | T1 | 6 | 6 | 0 |
| 66-11 | T1 | 7 | 7 | 0 |
| 66-18 | T1 | 6 | 6 | 0 |
| 66-16-7 | T2 | 5 | 5 | 0 |
| 66-16-8 | T2 | 8 | 8 | 0 |
| 67-16 | T1 | 6 | 6 | 0 |
| 67-17 | T1 | 5 | 5 | 0 |
| 225-3 | T1 | 7 | 7 | 0 |
| 226-4 | T1 | 8 | 8 | 0 |
| 67-16-10 | T2 | 5 | 5 | 0 |
| 67-16-12 | T2 | 3 | 3 | 0 |
| 67-16-2 | T2 | 2 | 2 | 0 |
| 67-16-8 | T2 | 7 | 7 | 0 |
| 67-16-9 | T2 | 9 | 9 | 0 |
| Total |  | 84 | 84 | 0 |

**Table S7.**
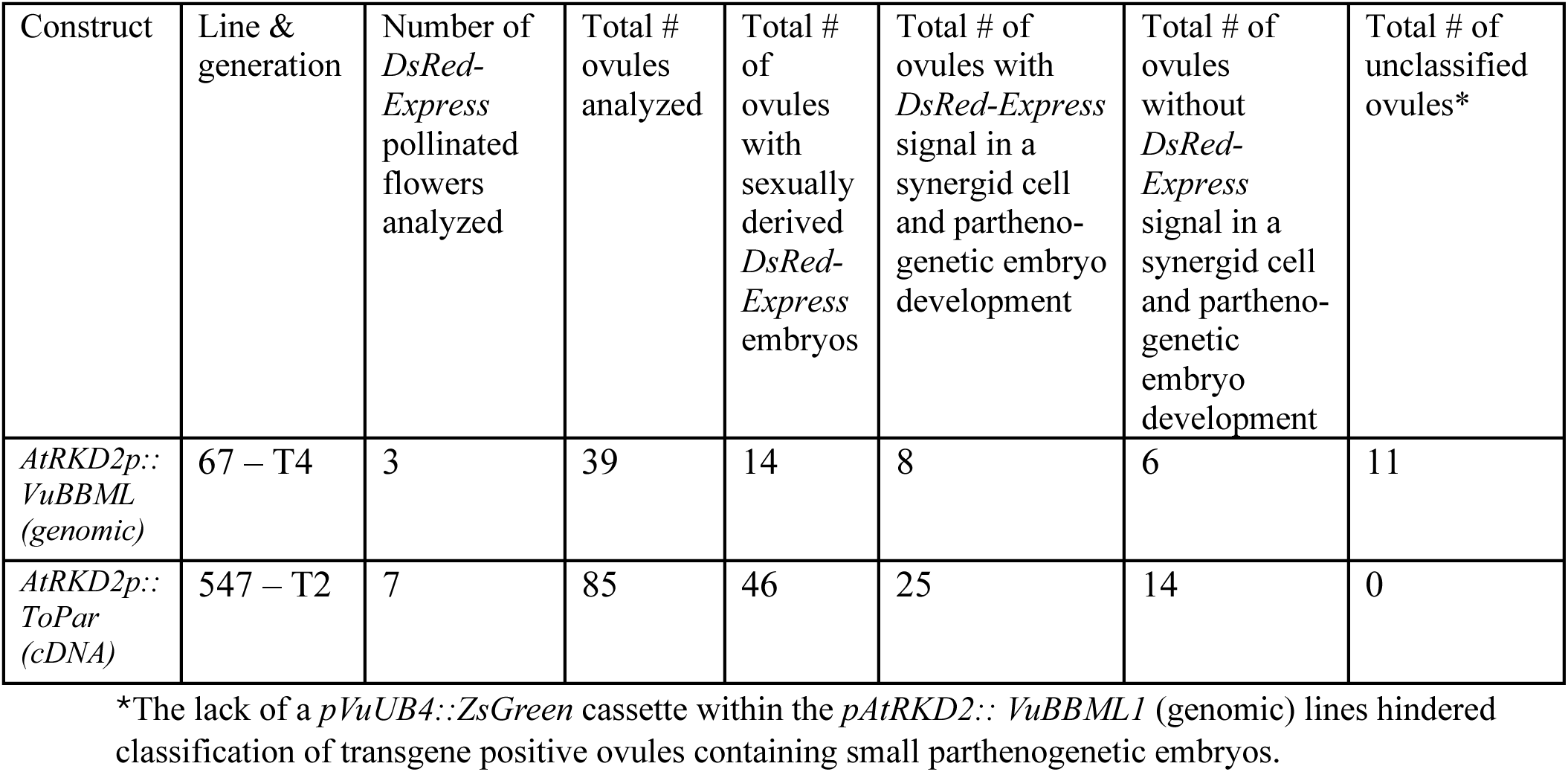
Quantification of pollen tube attraction in *pAtRKD2::VuBBML1* and *pAtRKD2::ToPAR* transformants.

**Table S8.** Quantification of seed set in mature pods of wild-type, T0 and T1 plants.

| Plant Id | # Pods analyzed | #Seeds harvested | # Normal size 2n seeds (%) | # Small size 1n seeds (%) | # Aborted seeds (%) | # Unfertilized ovules |
| --- | --- | --- | --- | --- | --- | --- |
| WT1 | 5 | 68 | 68 (100) | 0 | 0 | 12 |
| WT2 | 5 | 70 | 69 (98.5) | 0 | 1 (1.4) | 10 |
| **Total** | **10** | **138** | **137 (99.2)** | **0** | **1 (0.7)** | **22** |

*pAtDD45::VuBBML1*
|  | Plant Id | #Pods analyzed | #Seeds harvested | #Normal size 2n seeds (%) | #Small size 1n seeds (%) | # Aborted seeds (%) | #Unfertilized ovules |
| --- | --- | --- | --- | --- | --- | --- | --- |
| T0 individuals | sBBML1-764 | 3 | 15 | 11 (73.3) | 0 | 4 (26.7) | 35 |
|  | sBBML1-776 | 2 | 15 | 15 (100) | 0 | 0 | 15 |
|  | sBBML1-779 | 3 | 20 | 16 (80) | 0 | 4 (20) | 26 |
|  | **Total** | **8** | **50** | **42 (84)** | **0** | **8 (16)** | **76** |

*pAtDD25::VuBBML1-pAtDD45::VuBBML1*
|  | Plant Id | # Pods analyzed | # Seeds harvested | # Normal size 2n seeds (%) | # Small size 1n seeds (%) | # Aborted seeds (%) | # Unfertilized ovules |
| --- | --- | --- | --- | --- | --- | --- | --- |
| T0 individuals | dBBML1-690 | 5 | 43 | 40 (93) | 0 | 3 (7) | 37 |
|  | dBBML1-699 | 5 | 62 | 39 (62.9) | 13 (20.9) | 10 (16.1) | 18 |
|  | dBBML1-700 | 5 | 71 | 53 (74.6) | 11 (15.5) | 7 (9.8) | 9 |
|  | dBBML1-709 | 6 | 68 | 50 (73.5) | 7 (10.3) | 11 (16.1) | 28 |
|  | **Total** | **21** | **244** | **182 (74.5)** | **31 (12.7)** | **31 (12.7)** | **92** |
| T1 individuals | dBBML1690-1 | 5 | 70 | 55 (78.6) | 9 (12.6) | 6 (8.6) | 10 |
|  | dBBML1690-2 | 3 | 39 | 28 (71.8) | 3 (7.7) | 8 (20.5) | 9 |
|  | dBBML1699-1 | 5 | 74 | 50 (67.6) | 14 (18.9) | 10 (13.5) | 6 |
|  | dBBML1699-2 | 4 | 51 | 28 (54.9) | 9 (17.6) | 14 (27.4) | 13 |
|  | dBBML1700-1 | 3 | 30 | 16 (53.3) | 12 (40) | 2 (6.7) | 18 |
|  | dBBML1700-2 | 4 | 61 | 39 (63.9) | 14 (22.9) | 8 (13.1) | 3 |
|  | **Total** | **24** | **325** | **216 (66.4)** | **61 (18.8)** | **48 (14.8)** | **59** |

**Table S9.** Comparison of heterozygous SNP markers of a maternal *pAtDD25::VuBBML1-pAtDD45::VuBBML1* T0 plant dBBML699 and two of its haploid off-spring plants derived from self-pollination (ddVuBBML699-91 and dBBML699-98). The dBBML699-P1 and dBBML699-P2 columns depict the first and second allelic variant of 114 heterozygous SNP markers selected in the maternal plant to cover all 11 chromosomes. The dBBML699-91 and dBBML699-98 columns show the only allelic variant that was invariably found in two different haploid individuals for the same set of SNPs.

| Marker | Chr | dBBML699-P1 | dBBML-699-P2 | dBBML699-91 | dBBML699-98 |
| --- | --- | --- | --- | --- | --- |
| 1 | 1 | A | T | T | T |
| 2 | 1 | G | T | T | T |
| 3 | 1 | A | G | G | G |
| 4 | 1 | A | G | A | G |
| 5 | 1 | T | G | T | G |
| 6 | 1 | A | G | G | G |
| 7 | 1 | G | C | C | C |
| 8 | 1 | C | T | T | C |
| 9 | 1 | A | G | G | A |
| 10 | 2 | A | T | T | A |
| 11 | 2 | T | C | C | T |
| 12 | 2 | G | A | A | G |
| 13 | 2 | C | A | C | C |
| 14 | 2 | C | T | C | T |
| 15 | 2 | A | G | A | G |
| 16 | 2 | G | A | G | A |
| 17 | 2 | A | T | A | T |
| 18 | 2 | A | C | C | C |
| 19 | 2 | T | C | C | C |
| 20 | 2 | A | G | G | A |
| 21 | 2 | A | C | C | C |
| 22 | 2 | A | G | G | G |
| 23 | 2 | T | G | T | G |
| 24 | 2 | G | A | G | A |
| 25 | 2 | C | A | C | C |
| 26 | 2 | T | G | T | G |
| 27 | 2 | T | C | T | C |
| 28 | 3 | T | G | G | T |
| 29 | 3 | T | A | A | T |
| 30 | 3 | A | G | A | A |
| 31 | 3 | C | T | C | T |
| 32 | 3 | C | A | C | A |
| 33 | 3 | A | G | A | G |
| 34 | 3 | T | A | T | A |
| 35 | 3 | G | T | G | T |
| 36 | 4 | C | T | C | C |
| 37 | 4 | T | A | T | T |
| 38 | 4 | T | A | T | A |
| 39 | 4 | A | C | A | A |
| 40 | 4 | T | G | T | T |
| 41 | 4 | C | A | C | C |
| 42 | 4 | G | T | G | G |
| 43 | 5 | C | G | C | C |
| 44 | 5 | G | T | T | G |
| 45 | 5 | G | T | T | G |
| 46 | 5 | C | T | T | T |
| 47 | 5 | A | G | A | A |
| 48 | 6 | T | G | G | T |
| 49 | 6 | G | C | C | G |
| 50 | 6 | A | G | G | A |
| 51 | 6 | C | A | A | C |
| 52 | 6 | T | A | T | T |
| 53 | 6 | A | T | T | A |
| 54 | 6 | T | G | G | T |
| 55 | 6 | C | T | T | C |
| 56 | 6 | C | T | T | C |
| 57 | 7 | G | A | G | A |
| 58 | 7 | C | G | G | C |
| 59 | 7 | C | A | A | C |
| 60 | 7 | G | A | A | G |
| 61 | 7 | G | T | T | G |
| 62 | 7 | A | A | A | A |
| 63 | 7 | T | A | T | A |
| 64 | 7 | G | A | G | A |
| 65 | 7 | A | G | A | G |
| 66 | 7 | T | C | C | C |
| 67 | 7 | C | T | T | T |
| 68 | 8 | C | T | T | T |
| 69 | 8 | A | C | C | C |
| 70 | 8 | A | G | G | A |
| 71 | 8 | T | G | G | T |
| 72 | 8 | T | C | C | T |
| 73 | 8 | C | G | G | C |
| 74 | 8 | A | G | G | A |
| 75 | 8 | G | A | A | A |
| 76 | 8 | T | A | A | A |
| 77 | 8 | G | T | T | T |
| 78 | 8 | A | T | T | T |
| 79 | 8 | A | C | C | c |
| 80 | 9 | T | G | T | T |
| 81 | 9 | C | A | A | C |
| 82 | 9 | T | A | A | T |
| 83 | 9 | A | G | G | A |
| 84 | 9 | G | T | G | G |
| 85 | 9 | T | C | T | T |
| 86 | 9 | T | G | T | T |
| 87 | 9 | T | G | T | T |
| 88 | 9 | C | A | C | A |
| 89 | 9 | A | T | T | T |
| 90 | 9 | G | A | A | A |
| 91 | 9 | C | A | A | A |
| 92 | 9 | C | A | A | A |
| 93 | 10 | G | T | T | T |
| 94 | 10 | A | T | A | T |
| 95 | 10 | A | C | C | C |
| 96 | 10 | T | G | G | G |
| 97 | 10 | C | T | C | T |
| 98 | 10 | C | T | T | C |
| 99 | 10 | G | A | A | G |
| 100 | 10 | A | C | C | A |
| 101 | 10 | A | T | A | T |
| 102 | 10 | T | G | T | G |
| 103 | 11 | G | A | G | A |
| 104 | 11 | C | A | C | A |
| 105 | 11 | G | T | T | T |
| 106 | 11 | T | G | T | G |
| 107 | 11 | T | G | T | G |
| 108 | 11 | G | C | G | G |
| 109 | 11 | G | T | G | G |
| 110 | 11 | G | T | G | G |
| 111 | 11 | T | C | C | C |
| 112 | 11 | C | A | C | C |
| 113 | 11 | A | G | A | A |
| 114 | 11 | A | C | A | A |

**Table S10.**
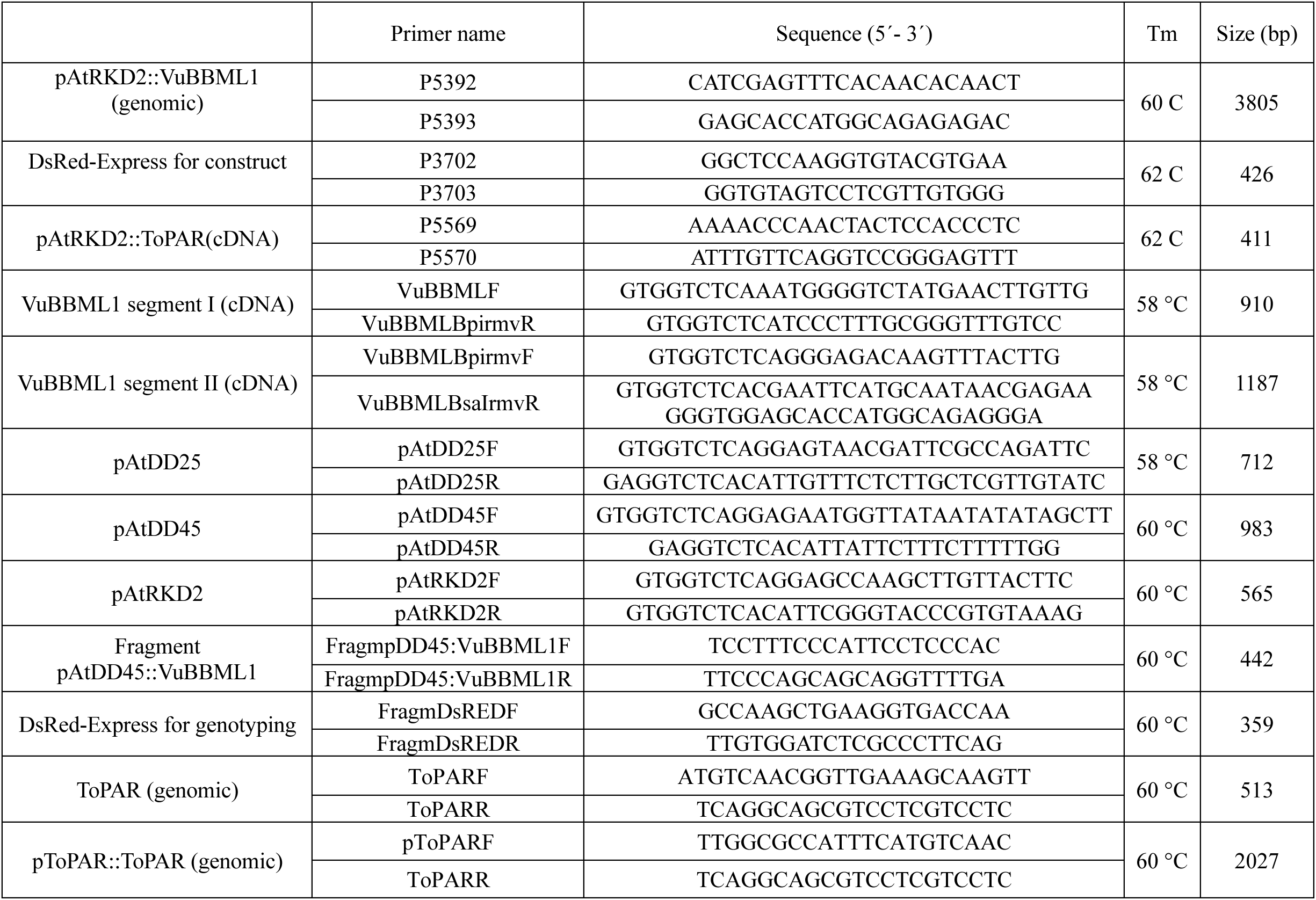
Primers used in this study.

